# Phage-like particle vaccines are highly immunogenic and protect against pathogenic coronavirus infection and disease

**DOI:** 10.1101/2021.11.08.467648

**Authors:** Bennett J. Davenport, Alexis Catala, Stuart M. Weston, Robert M. Johnson, Jeremy Ardunay, Holly L. Hammond, Carly Dillen, Matthew B. Frieman, Carlos E. Catalano, Thomas E. Morrison

## Abstract

The response by vaccine developers to the COVID-19 pandemic has been extraordinary with effective vaccines authorized for emergency use in the U.S. within one year of the appearance of the first COVID-19 cases. However, the emergence of SARS-CoV-2 variants and obstacles with the global rollout of new vaccines highlight the need for platforms that are amenable to rapid tuning and stable formulation to facilitate the logistics of vaccine delivery worldwide. We developed a “designer nanoparticle” platform using phage-like particles (PLPs) derived from bacteriophage lambda for multivalent display of antigens in rigorously defined ratios. Here, we engineered PLPs that display the receptor binding domain (RBD) protein from SARS-CoV-2 and MERS-CoV, alone (RBD_SARS_-PLPs, RBD_MERS_-PLPs) and in combination (hCoV-RBD PLPs). Functionalized particles possess physiochemical properties compatible with pharmaceutical standards and retain antigenicity. Following primary immunization, BALB/c mice immunized with RBD_SARS_- or RBD_MERS_-PLPs display serum RBD-specific IgG endpoint and live virus neutralization titers that, in the case of SARS-CoV-2, were comparable to those detected in convalescent plasma from infected patients. Further, these antibody levels remain elevated up to 6 months post-prime. In dose response studies, immunization with as little as one microgram of RBD_SARS_-PLPs elicited robust neutralizing antibody responses. Finally, animals immunized with RBD_SARS_-PLPs, RBD_MERS_-PLPs, and hCoV-RBD PLPs were protected against SARS-CoV-2 and/or MERS-CoV lung infection and disease. Collectively, these data suggest that the designer PLP system provides a platform for facile and rapid generation of single and multi-target vaccines.

## INTRODUCTION

Severe acute respiratory syndrome coronavirus 2 (SARS-CoV-2), a positive-sense, single-stranded RNA virus, was first isolated in late 2019 from patients with severe respiratory illness in Wuhan, China (*1, 2*). This betacoronavirus is related to two other coronaviruses that are highly pathogenic for humans – SARS-CoV and Middle East respiratory syndrome coronavirus (MERS-CoV) (*3*). Although SARS-CoV no longer circulates in the human population, SARS-CoV-2 infection is the cause of the global pandemic of coronavirus disease 2019 (COVID-19), which can progress to acute respiratory distress syndrome (ARDS) and death (*4*). The elderly, immunocompromised, and those with certain co-morbidities (e.g., obesity, diabetes, and hypertension) are at greatest risk of severe COVID-19 (*5*). In addition, MERS-CoV remains in circulation and has infected over 2,500 people with lethal disease in ∼834 (∼34% case fatality rate)(*6*). The virus has spread to 28 countries since emerging in the Kingdom of Saudi Arabia in 2012 (*6, 7*), and large outbreaks of human-to-human transmission have occurred in Riyadh and Jeddah in 2014 and in South Korea in 2015 (*8, 9*).

The RNA genome of betacoronaviruses is approximately 30,000 nucleotides in length (*10*). The 5′ two-thirds encode nonstructural proteins that catalyze genome replication and viral RNA synthesis, whereas the 3′ one-third of the genome encodes the viral structural proteins, including nucleoprotein and the spike (S), envelope, and membrane proteins. The S protein of coronaviruses forms homotrimeric spikes on the virion (**Fig 1A**), which engages cell surface attachment factors and entry receptors (*11*). Protein cleavage occurs sequentially during the entry process to yield S1 and S2 fragments and is then followed by further processing of S2 to yield a smaller S2′ protein (*12*). The S1 fragment includes the receptor binding domain (RBD), the predominant target of potently neutralizing monoclonal antibodies, while the S2 fragment promotes membrane fusion (*13-22*).

**Figure 1.**
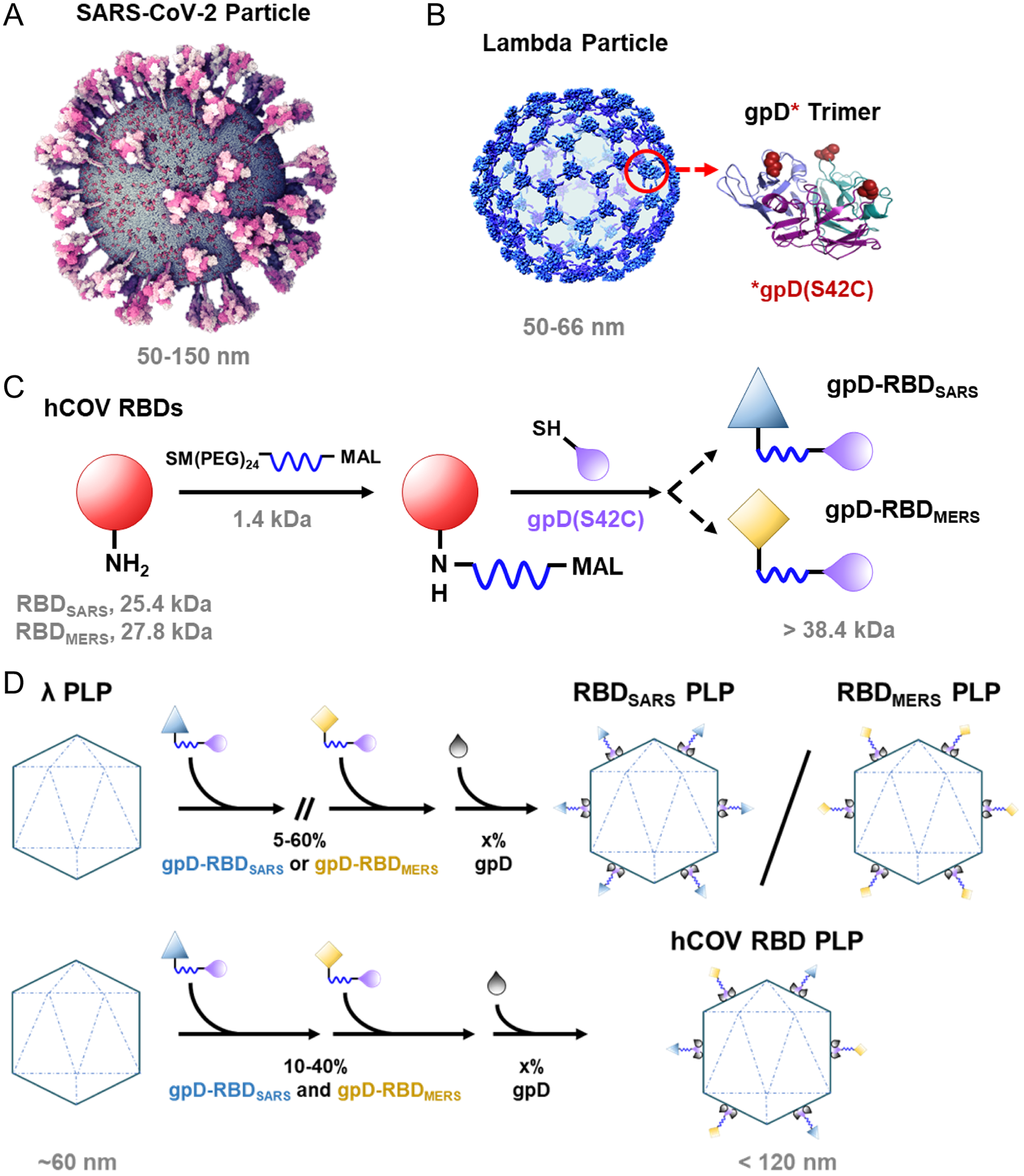
Design and construction of hCoV RBD decorated particles. (**A**) Atomistic model of the SARS-CoV-2 virion with homotrimeric spikes of the S protein shown in shades of pink (Nanographics GmbH). (**B**) Cryo-EM reconstruction of the phage lambda capsid with only the density from the trimeric spikes of the decoration protein, gpD, displayed (Lander et al., 2008). Shown to the right in cartoon representation is the crystal structure of an isolated gpD trimer (PDB: 1C5E) modified to contain the serine to cysteine mutation (red spheres) of the gpD variant, gpD(S42C). (**C**) Reaction schematic for the chemical crosslinking of human coronavirus RBD proteins (hCOV RBDs) to gpD(S42C) *via* lysine amide coupling and thiol-maleimide chemistry. The products (gpD-RBD_SARS_, gpD-RBD_MERS_) contain a PEG crosslinker, SM(PEG)_24_, with a spacer arm length of 95.2 Å. (**D**) PLPs are decorated with RBDs at the desired surface densities either independently (top) or in combination (bottom) with remaining gpD-binding sites filled using wildtype gpD such that all 420 binding sites are occupied.

A number of vaccine candidates against coronaviruses, targeting the S protein, have been developed, including DNA plasmid, lipid nanoparticle encapsulated mRNA, inactivated virion, and viral-vectored vaccines (*23, 24*). For instance, currently approved RNA-based vaccines that target the S protein of SARS-CoV-2 are safe and highly effective (*25, 26*); however, they suffer from multiple-dose requirements and require strict “cold-chain” conditions that limit vaccine distribution and pose constraints on the infrastructure and healthcare system. Conversely, adenovirus vector vaccines against SARS-CoV-2 can have the advantage of single-dose administration (*27*), but they are less effective than RNA vaccines and can lead to serious side effects, such as labored breathing, transient neutropenia and lymphopenia, liver damage, and, in rare cases, neuropathies like Bell’s palsy and transverse myelitis (*28-30*). Furthermore, the success of platforms derived from eukaryotic viruses (e.g., adenoviruses) has been tempered by safety concerns related to pathogenicity, immunogenicity, and toxicity (*31*). Thus, there continues to be a need for multifaceted, rapidly tunable vaccine platforms that can respond to novel emerging threats, including natural variants, that provide long-term protection, and that are amenable to industrial strategies to afford thermostable, single-shot formulations for efficient distribution in resource-limited settings.

Bacteriophage systems have been adapted as therapeutic and diagnostic (theranostic) platforms. In these cases, phage-like particles (PLPs) derived from capsid proteins are utilized as a scaffold for genetic and chemical modification (*32, 33*). For example, filamentous phages, such as M13 and fd, and icosahedral phages, including P22, T4, AP205, MS2, and lambda, have been used to display imaging reagents (*34-36*), synthetic polymers (*37-39*), bioactive peptides and proteins (*39-46*). PLPs have also been employed as vaccine platforms to display antigens including from *Y. pestis* (*42*) and *P. falciparum* (*43*). In these cases, the PLP decoration ligands are engineered using genetic and/or chemical modification of the capsid-associated proteins.

The Catalano laboratory has developed a “designer nanoparticle” platform adapted from phage lambda (*47*). Co-expression of the major capsid and “scaffolding” proteins in *Escherichia coli* affords icosahedral PLP shells that can be isolated in high yield. These can then be decorated *in vitro* with the lambda decoration protein (gpD) such that a single particle will display 140 trimeric spikes of gpD (420 total copies) projecting from its surface (*48, 49*). The decoration protein can be modified genetically to display heterologous peptides and proteins that can be displayed on the PLP surface (*39, 50, 51*). Additionally, we have engineered a mutant gpD protein that contains a Ser42->Cys mutation (gpD(S42C)), which enables site-specific chemical modification of the decoration protein using maleimide chemistry (**Fig 1B**) (*39*). Employing our *in vitro* system, PLPs can be independently or simultaneously decorated with genetically and chemically modified decoration proteins in rigorously defined surface densities (*39, 46*). Notably, the lambda PLP platform has several advantages and unique features that can be harnessed for vaccine development. For example, (i) PLPs can be decorated under defined *in vitro* conditions with biological and synthetic molecules in varying surface densities using genetic and chemical modification strategies; (ii) modification of the particle surface is fast and can be tuned in a user defined manner, thereby streamlining the formulation process; (iii) intact phage and PLP shells are natural adjuvants capable of stimulating the innate immune response (*52, 53*); (iv) the ability to display antigens in high density can aid in regulating effector function (*54*); (v) decorated particles are monodisperse, stable, and possess physiochemical properties amenable for pharmaceutical formulation and therapeutic applications (*46*). Therefore, the lambda system described here allows for substantial flexibility and rigorous control in vaccine design and development, setting the stage for the construction of an “all-in-one” vaccine platform.

In this study, we report the development of a monovalent lambda PLP-based vaccine against SARS-CoV-2 (RBD_SARS_-PLP) or MERS-CoV (RBD_MERS_-PLP) via decoration with the spike RBD proteins from either virus. Additionally, we engineered bivalent PLPs that co-display spike RBD proteins from both viruses (hCoV-RBDs-PLP) to serve as a bivalent vaccine candidate. Intramuscular administration of RBD_SARS_-PLPs, RBD_MERS_-PLPs, and hCoV-RBDs PLPs in mice induce robust and durable humoral immune responses, including the production of neutralizing antibodies. Moreover, immunization also protected mice against lung infection, inflammation, and pathology after virulent SARS-CoV-2 and MERS-CoV challenge.

## RESULTS

### Design and construction of particles decorated with human coronavirus spike RBD proteins

The lambda designer PLP platform was adapted to display CoV spike RBD proteins using methods established in the Catalano laboratory (*39, 46*). Purified recombinant spike RBD proteins derived from the SARS-CoV-2 and MERS-CoV isolates Wuhan-Hu-1 and EMC/2012 are referred to as RBD_SARS_ and RBD_MERS_, respectively, or collectively as hCoV-RBDs. These proteins were crosslinked to the lambda decoration protein mutant, gpD(S42C), following a two-step procedure. First, solvent accessible lysine residues in the hCoV-RBDs were modified with the *N*-hydroxysuccinimide ester groups of the SM(PEG)_24_ heterobifunctional crosslinker, as depicted in **Fig 1C**; SDS-PAGE analysis reveals an apparent single predominant product for both RBD_SARS_ and RBD_MERS_ (**Fig 2A,D**, respectively, *lanes 4*). Next, the maleimide groups of the crosslinker were reacted with the sole cysteine residue of gpD(S42C) to afford the gpD-RBD_SARS_ and gpD-RBD_MERS_ constructs (**Fig 1C**); SDS-PAGE analysis reveals multiple products in each case (**Fig 2A,D**, *lanes 5*). The reaction mixtures were fractionated by size exclusion chromatography (SEC) (**Fig 2B,E**), and the fraction containing a single predominant product for each hCoV RBD was isolated and then concentrated for use in all subsequent experiments (*fraction 32*, **Fig 2C,F**).

**Figure 2.**
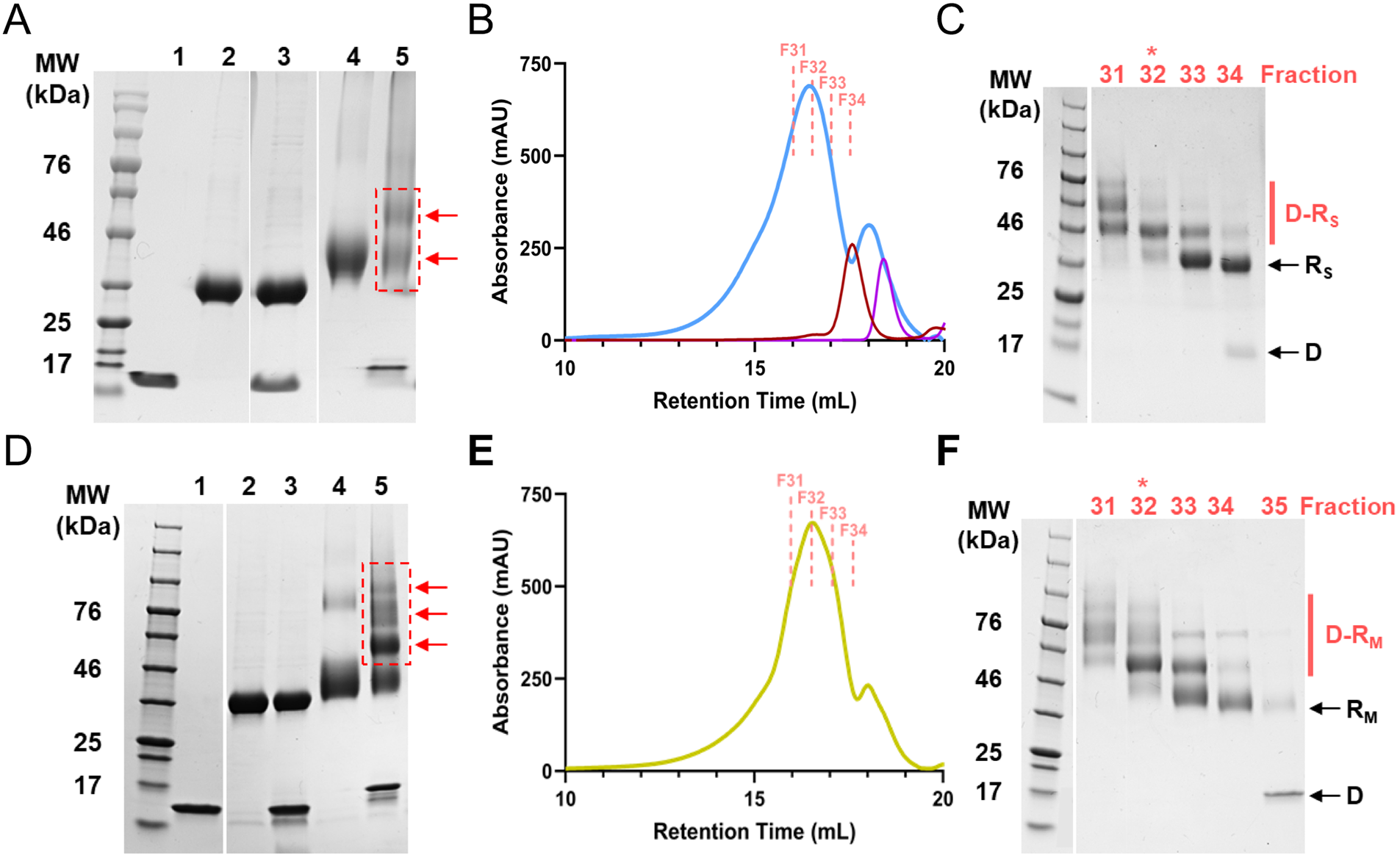
Construction of hCoV RBDs. (**A**) Proteins at various steps in the chemical modification of gpD(S42C) for conjugation to RBD_SARS_ were fractionated by SDS-PAGE and visualized by Coomassie Blue staining. Lanes: (1) gpD(S42C); (2) RBD_SARS_; (3) gpD(S42C) and RBD_SARS_ in the absence of crosslinker; (4) RBD_SARS_ modified with the crosslinker; (5) gpD-RBD_SARS_ crosslinked proteins (red arrows). (**B**) SEC chromatogram of the gpD-RBD_SARS_ reaction mixture (blue); collected fractions numbers are indicated at the top. SEC chromatograms of unmodified RBD_SARS_ (red) and unmodified gpD(S42C) (purple) are shown for comparison. (**C**) SDS-PAGE of fractions eluting from the SEC column in panel B. The migration of purified gpD-RBD_SARS_, unmodified RBD_SARS_, and gpD(S42C) are denoted as D-R_S_, R_S_, and D, respectively. Unless otherwise stated, gpD-RBD_SARS_ fraction 32 (red star) was used for all subsequent experiments. (**D**) Proteins at various steps in the chemical modification of gpD(S42C) for conjugation to RBD_MERS_ were fractionated by SDS-PAGE and visualized by Coomassie Blue staining. Lanes: (1) gpD(S42C); (2) RBD_MERS_; (3) gpD(S42C) and RBD_MERS_ in the absence of crosslinker; (4) RBD_MERS_ modified with the crosslinker; (5) gpD-RBD_MERS_ crosslinked proteins (red arrows). (**E**) SEC chromatogram of the gpD-RBD_MERS_ reaction mixture (yellow); collected fractions are indicated at the top. (**F**) SDS-PAGE of fractions eluting from the SEC column in panel E. The migration of purified gpD-RBD_MERS_, unmodified RBD_MERS_, and gpD(S42C) are denoted as D-R_M_, R_M_, and D, respectively. Unless otherwise stated, gpD-RBD_MERS_fraction 32 (red star) was used for all subsequent experiments.

The purified gpD-RBD constructs were the used to decorate PLPs, alone and in combination at defined surface densities, as outlined in **Fig 1D**. Complete particle decoration requires 420 copies of the decoration protein that assemble as trimeric spikes, projecting from the shell surface (see **Fig 1B**); therefore, we define the number of gpD-binding sites occupied as a surface density percentage. For instance, 60% RBD_SARS_-PLPs denotes particles that are each decorated with 252 copies of gpD-RBD_SARS_ with the remaining sites filled with wildtype (WT) gpD (168 copies). Similarly, 40% hCoV-RBD PLPs are particles that are each decorated with 168 copies of gpD-RBD_SARS_, 168 copies of gpD-RBD_MERS_, and 84 copies of WT gpD. Analysis of the reaction mixtures by agarose gel electrophoresis (AGE) demonstrated that particles can be decorated in a defined manner with either and both gpD-RBD constructs (**Fig 3A-C**).

**Figure 3.**
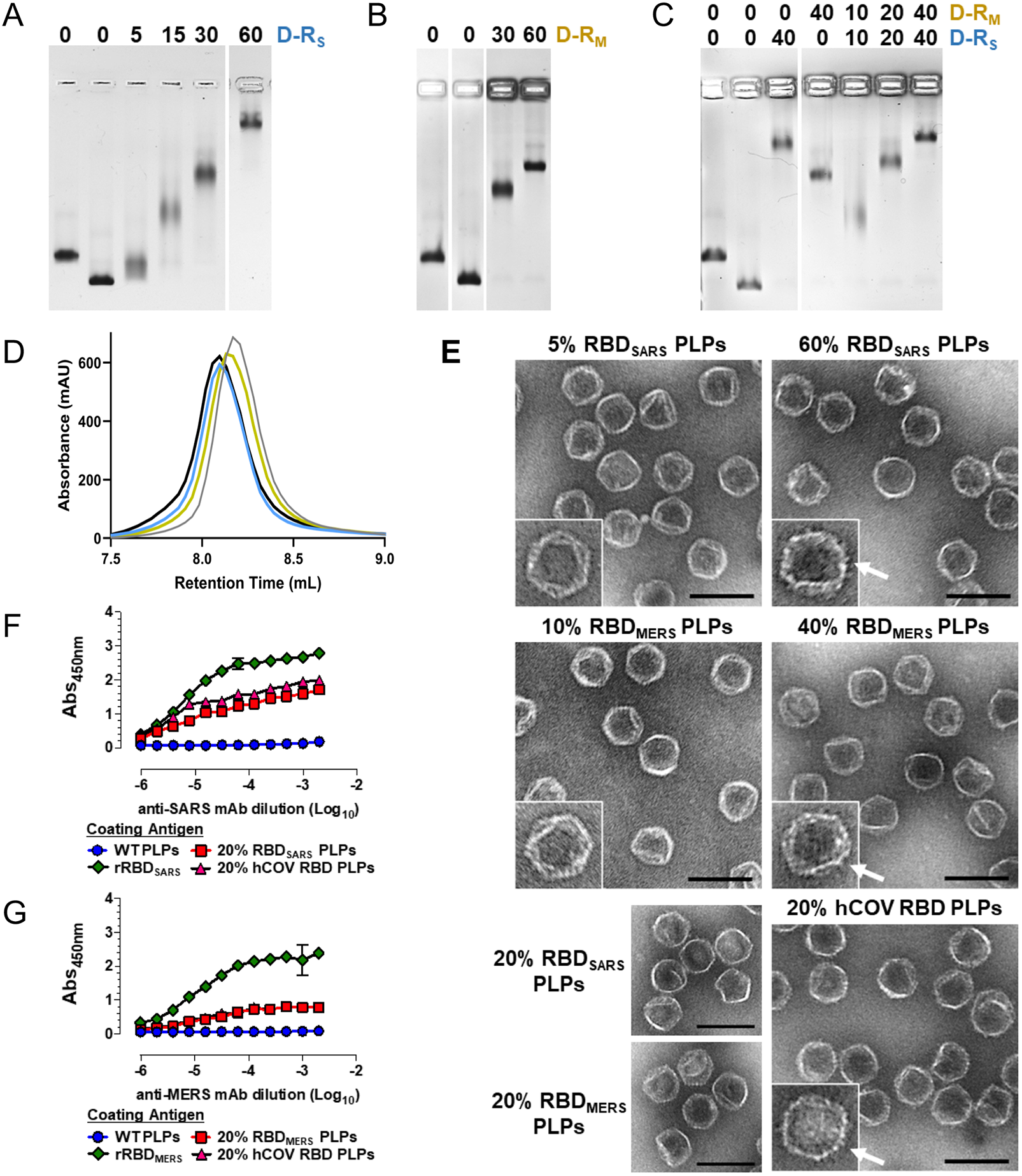
PLP decoration with hCoV RBDs. PLPs were decorated with (**A**) gpD-RBD_SARS_(D-R_S_, blue), (**B**) gpD-RBD_MERS_ (D-R_M_, yellow) or (**C**) with both constructs at the indicated surface densities as outline in Fig 1D and the reaction mixtures were fractionated by AGE. In all gels, naked PLPs (no decoration protein, *lanes 1*) and WT PLPs (decorated with 100% wildtype gpD (grey), *lanes 2*) served as controls for the decoration reactions. (**D**) Representative SEC chromatograms showing WT PLPs (grey; retention time (RT), 8.2 mL), 20% RBD_SARS_ PLPs (blue; RT, 8.1 mL), 20% RBD_MERS_ PLPs (yellow; RT, 8.1 mL), and 20% hCOV RBD PLPs (black; RT, 8.1 mL). (**E**) Electron micrographs of PLPs decorated with hCOV RBDs (magnification, 98,000x; scale bars, 100 nm). White arrows indicate the surface density attributed to the RBD proteins projecting from the PLP surface. (**F-G**) Antigenicity of the purified hCoV-PLPs. ELISA plates were coated with 0.2 μg WT PLPs, 20% RBD_SARS_ PLPs, 20% RBD_MERS_ PLPs, 20% hCoV PLPs, and recombinant RBD protein as indicated. Plates were then probed with 2-fold serial dilutions of a detection antibody, and signals were developed following standard ELISA protocols. For (**F**), recombinant SARS-CoV-2 RBD protein is denoted as rRBD_SARS_, and plates were probed with an anti-SARS-CoV-2 RBD monoclonal antibody. For (**G**), recombinant MERS-CoV RBD protein is denoted as rRBD_MERS_, and plates were probed with an anti-MERS-CoV RBD detection antibody.

Decorated particles were purified by SEC (**Fig 3D**) and characterized by multiple approaches. Electron microscopy (EM) confirmed that decorated PLPs retain icosahedral symmetry and are well dispersed. Close inspection of the micrographs revealed that particles decorated with RBD densities greater than 20% are characterized by surface projections whose number increases with increasing surface decoration (**Fig 3E**); we attribute these projections to RBD proteins extending from the particle surface. Physiochemical characterization of PLP preparations by dynamic and electrophoretic light scattering analyses reveal that the mean hydrodynamic diameter of particles decorated with RBDs also increases as a function of RBD surface density and have an overall surface charge that ranges from -12 to -29 mV (**Table 1**). The polydispersity index (PDI) of the decorated particles ranges from (0.156 ± 0.05) to (0.285 ± 0.09). In sum, these data indicate that these particle preparations have a high degree of homogeneity and are acceptable by pharmaceutical standards.

**TABLE 1.** CHARACTERIZATION OF PLP PREPARATIONS.

| Sample | Zetasizer* |  |  |  |  |
| --- | --- | --- | --- | --- | --- |
|  | Z-Ave<br>(d.nm) | PDI | Intensity<br>PSD (d.nm) | Volume<br>PSD (d.nm) | ZP (mV) |
| Naked PLPs | 66.2 ± 1.4 | 0.073 ± 0.01 | 71.8 ± 2.1 | 58.6 ± 1.1 | -25.1 ± 0.8 |
| WT PLPs | 73.8 ± 1.3 | 0.081 ± 0.04 | 80.1 ± 3.0 | 65.6 ± 1.6 | -17.4 ± 2.4 |
| 5% RBD<sub>SARS</sub> PLPs | 82.4 ± 3.0 | 0.156 ± 0.05 | 90.8 ± 2.4 | 68.5 ± 0.7 | -18.8 ± 2.4 |
| 20% RBD<sub>SARS</sub> PLPs | 111.5 ± 6.5 | 0.285 ± 0.09 | 121.3 ± 6.0 | 86.8 ± 5.2 | -28.5 ± 1.6 |
| 60% RBD<sub>SARS</sub> PLPs | 110.7 ± 7.1 | 0.223 ± 0.06 | 131.9 ± 17 | 94.4 ± 6.7 | -11.5 ± 2.6 |
| 10% RBD<sub>MERS</sub> PLPs | 84.3 ± 3.3 | 0.164 ± 0.05 | 89.7 ± 1.5 | 70.7 ± 0.6 | -11.8 ± 1.4 |
| 20% RBD<sub>MERS</sub> PLPs | 94.9 ± 3.4 | 0.212 ± 0.06 | 101.7 ± 4.2 | 76.6 ± 1.4 | -22.2 ± 2.1 |
| 40% RBD<sub>MERS</sub> PLPs | 93.1 ± 3.3 | 0.176 ± 0.04 | 102.3 ± 4.8 | 77.7 ± 1.7 | -15.2 ± 2.1 |
| 20% hCOV RBD PLPs | 115.3 ± 16 | 0.277 ± 0.11 | 112.9 ± 9.1 | 85.7 ± 5.2 | -23.3 ± 2.8 |

\*PLP preparations were diluted in 10 mM HEPES [pH 7.4], and measurements acquired in quadruplicate at 25°C. Values shown are mean ± standard deviation.
Particle sizing data is based on particle diameter (d.nm) as measured by dynamic light scattering from two to three independent analyses. These are reported as Z-average (Z-Ave; weighted mean hydrodynamic size), polydispersity index (PDI; measure of sample heterogeneity) and particle size distributions (PSD; intensity, volume).
Particle charge data is based on overall surface charge (Zeta potential, ZP) as measured by electrophoretic light scattering.

Having confirmed that PLPs can be decorated with varying surface densities of hCoV RBDs, we sought to determine whether the purified gpD-RBD constructs retain their antigenic characteristics following particle decoration. Thus, the antigenicity of RBD_SARS_-PLPs, RBD_MERS_-PLPs, and hCoV-RBD-PLPs was assessed using enzyme immunoassays. The data presented in **Figs 3F** and **3G** show that PLPs decorated with 20% gpD-RBD_SARS_ and/or 20% gpD-RBD_MERS_ retain antigenic properties, whereas no reactivity was noted for WT PLPs.

### Particles decorated with gpD-RBD_SARS_ are immunogenic following one or two immunizations

To assess the immunogenicity of RBD_SARS_ PLPs, 6-week-old BALB/c mice were immunized by intramuscular (i.m.) inoculation with 10 ug of WT PLPs (control) or 60% RBD_SARS_-PLPs, followed by administration of a booster dose three weeks later (**Fig 4A**). Serum samples were collected 14 days after the primary or booster immunization (day 35 post-prime) and again at days 63 and 174 post-prime. IgG responses against purified RBD protein were evaluated by ELISA (**Fig 4B**). After a single immunization, 60% RBD_SARS_-PLPs induced RBD_SARS_-specific IgG titers comparable to those detected in convalescent samples from SARS-CoV-2 recovered patients (**Fig 4B,C**). At 14 days post-boost (day 35), these responses were elevated 16-fold (*P* < 0.001). Importantly, mice immunized with 60% RBD_SARS_-PLPs maintained high levels of RBD_SARS_-specific IgG out to day 174 post-prime (**Fig 4B**), indicating that these particles elicit durable humoral immune responses. In contrast, RBD_SARS_-specific IgG was undetectable in mice immunized with WT PLPs at all time points evaluated (**Fig 4B**). Furthermore, IgG subclass analysis at 14 days post-prime and boost revealed that 60% RBD_SARS_-PLPs elicit high levels of RBD_SARS_-specific IgG1, IgG2a, and IgG2b (**Fig 4D**).

**Figure 4.**
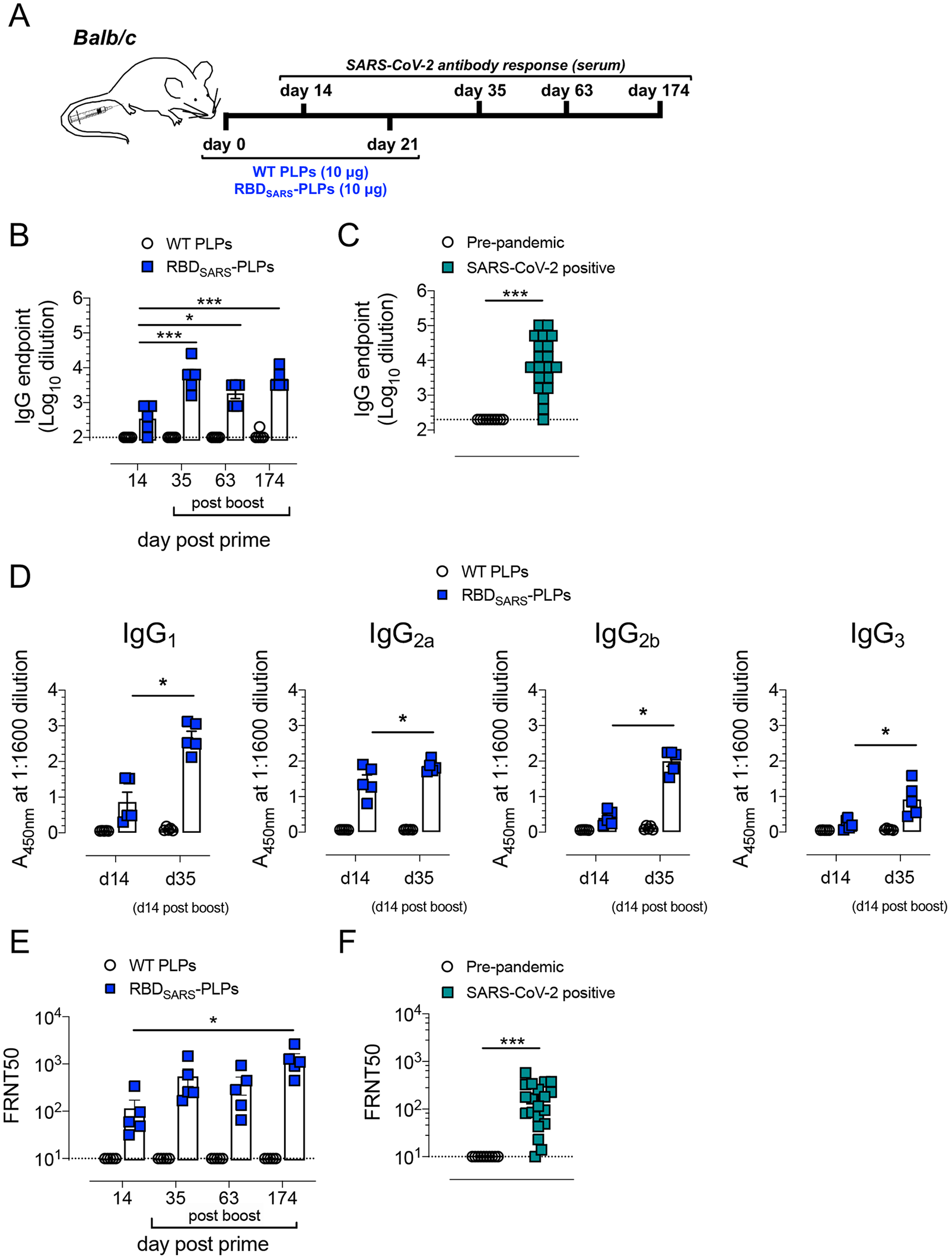
RBD_SARS_-PLPs are immunogenic. (**A**) Schematic of immunization protocol and sample collection. (**B**) WT BALB/c mice (n = 5 mice/group) were immunized with 10 μg of WT PLPs (control) or 60% RBD_SARS_-PLPs by intramuscular (i.m.) injection on days 0 and 21. Animals were bled on days 14, 35, 63, and 174 and RBD_SARS_-specific IgG endpoint titers were determined by ELISA. *P* values were determined by one-way ANOVA with Tukey’s multiple comparisons test. ^*^*P*<0.05, ^***^*P*<0.001. (**C**) RBD_SARS_-specific IgG ELISA endpoint titers in plasma from pre-pandemic controls (n = 10) and convalescent SARS-CoV-2 PCR positive patients (n = 20). **(D)** RBD_SARS_-specific IgG subclass responses on days 14 and 35 were determined by ELISA. *P* values were determined by unpaired student’s t test. ^*^*P*<0.05. (**E-F**) SARS-CoV-2 neutralizing activity was determined by a focus reduction neutralization test (FRNT). (**E**) The dilution of serum that inhibited 50% of infectivity (FRNT50 titer) was calculated for each sample by nonlinear regression analysis, as described in Materials and Methods. *P* values were determined by one-way ANOVA with Tukey’s multiple comparisons test. ^*^*P*<0.05, ^***^*P*<0.001. (**F**) SARS-CoV-2 FRNT50 titers in plasma from pre-pandemic controls (n = 10) and convalescent SARS-CoV-2 PCR positive patients (n = 20).

Additionally, we evaluated serum from immunized animals for the capacity to neutralize SARS-CoV-2 infection using a live virus focus-reduction neutralization test (FRNT). Serum from mice immunized with WT PLPs did not display neutralizing activity (**Fig S1**), whereas serum from mice immunized with 60% RBD_SARS_-PLPs 14 days post-prime neutralized SARS-CoV-2 infection comparable to the neutralizing activity detected in convalescent samples from SARS-CoV-2 recovered patients (**Fig 4E-F**). Neutralizing activity was enhanced 4.7-fold following the booster immunization (*P* < 0.05), and high levels of SARS-CoV-2 neutralizing antibodies were maintained out to day 174 post-prime (**Fig 4E and Fig S1**).

### Vaccination with RBD_SARS_-PLPs protects from virulent SARS-CoV-2 challenge

We evaluated the protective activity of particles decorated with gpD-RBD_SARS_ using a recently developed mouse-adapted strain of SARS-CoV-2 (SARS-CoV-2 MA10), which productively replicates in the mouse lung and results in clinical manifestations of disease consistent with severe COVID-19 in humans (*55*). Six-week-old BALB/c mice were immunized i.m. with WT PLPs or 60% RBD_SARS_-PLPs. At 184 days post-prime, mice were challenged intranasally with 10^4^ plaque-forming units (PFUs) of SARS-CoV-2 MA10 (**Fig 5**). Mice immunized with WT PLPs rapidly lost weight following SARS-CoV-2 infection, whereas no weight loss was observed in mice immunized with 60% RBD_SARS_-PLPs (**Fig 5A**). At 4 days post-infection (dpi), mice were euthanized, and lungs were collected for viral burden analysis and histopathology. While high levels of infectious virus were detected in the lungs of mice immunized with WT PLPs, infectious virus was undetectable in the lungs of mice immunized with 60% RBD_SARS_-PLPs, as determined by plaque assay (**Fig 5B**). Moreover, greatly reduced levels of genomic (**Fig 5C**) and subgenomic (**Fig 5D**) viral RNA were detected in the lungs of mice vaccinated with 60% RBD_SARS_-PLPs, as compared to those immunized with WT PLPs. Collectively, these data indicate that i.m. immunization with RBD_SARS_-PLPs elicits durable protection against SARS-CoV-2 lung infection.

**Figure 5.**
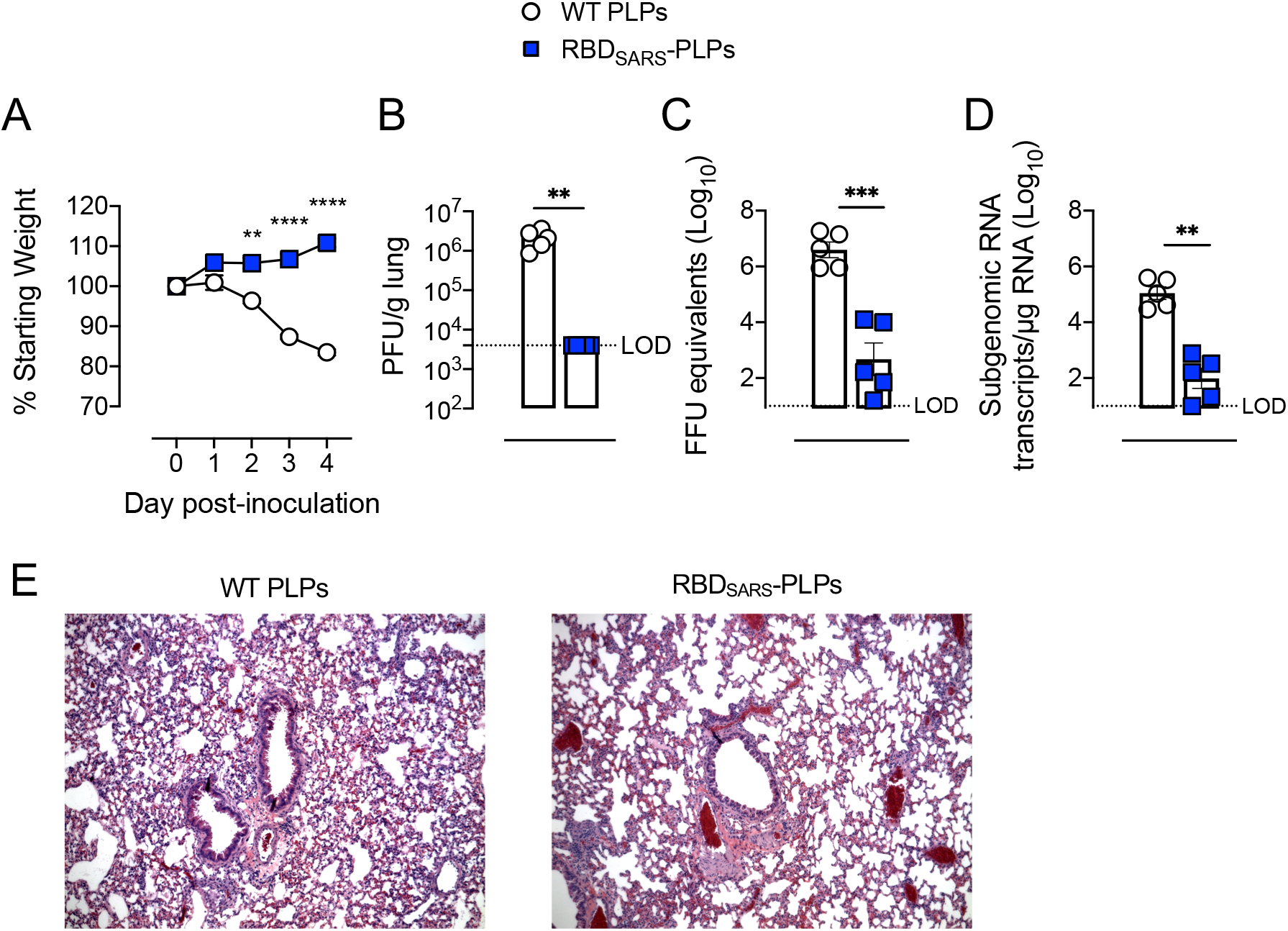
Immunization with RBD_SARS_-PLPs protects against SARS-CoV-2 infection and disease. (**A**) WT BALB/c mice (n = 5/group) were immunized with 10 μg of WT PLPs 60% RBD_SARS_-PLPs by intramuscular (i.m.) injection on days 0 and 21. At day 184, mice were challenged intranasally (i.n.) with 10^4^ PFU of mouse-adapted SARS-CoV-2 MA10. Mice were monitored daily for weight changes. *P* values were determined by two-way ANOVA with Tukey’s multiple comparisons test. ^**^*P*<0.01, ^****^*P*<0.0001. (**B-D**) At 4 dpi, viral burden in the lung was quantified by plaque assay (**B**), RT-qPCR for viral genomic RNA (**C**), and RT-qPCR for N subgenomic RNA (**D**). (**E**) Histopathology in lung tissue sections was evaluated by hematoxylin and eosin staining. *P* values were determined by Mann-Whitney test in **B, D**; or by unpaired student’s t-test in **C**. ^**^*P*<0.01, ^***^*P*<0.001.

The effect of immunization with RBD_SARS_-PLPs on lung inflammation and disease also was assessed by analyzing lung tissue for histopathological changes. Mice immunized with WT PLPs and challenged with SARS-CoV-2 MA10 had an abundant accumulation of immune cells in perivascular and alveolar locations, vascular congestion, and interstitial edema (**Fig 5E**). Conversely, immunization with 60% RBD_SARS_-PLPs resulted in a marked reduction of lung-associated histopathological changes (**Fig 5E**), indicative of protection against SARS-CoV-2 induced lung inflammation and injury.

### Low doses of RBD_SARS_-PLPs remain immunogenic and protect from virulent SARS-CoV-2 challenge

We next evaluated the immunogenicity and protective efficacy of de-escalating doses of particles decorated with gpD-RBD_SARS_. Six-week-old BALB/c mice were immunized by i.m. injection with 2.5, 1.0, or 0.25 μg of 60% RBD_SARS_-PLPs and received a booster dose of the same amount three weeks later. Control mice received two i.m. injections of 2.5 μg of WT PLPs, following the same immunization schedule. Serum samples were collected on days 14, 35 (14 days post-boost), 63, and 84 post-prime, and IgG responses against purified RBD protein were evaluated by ELISA. At 14 days, little response was observed in mice immunized with a single injection of 0.25 μg of 60% RBD_SARS_-PLPs, and no response was detected in control mice (**Fig 6A**). In contrast, mice immunized with 2.5 or 1.0 μg of 60% RBD_SARS_-PLPs had detectable levels of RBD_SARS_-specific IgG which were comparable to mice immunized with a 10 μg dose (compare **Figs 4B** and **6A**), and to RBD_SARS_-specific IgG detected in convalescent samples from SARS-CoV-2 recovered patients (**Fig 4C**). At 14 days post-boost (day 35 post-prime), RBD_SARS_-specific IgG responses were elevated in mice that received all doses of 60% RBD_SARS_-PLPs, although lower responses were observed for mice immunized with 0.25 μg (**Fig 6A**). Additionally, mice immunized with all doses maintained high levels of RBD_SARS_-specific IgG up to 84 days post-prime (**Fig 6A**).

**Figure 6.**
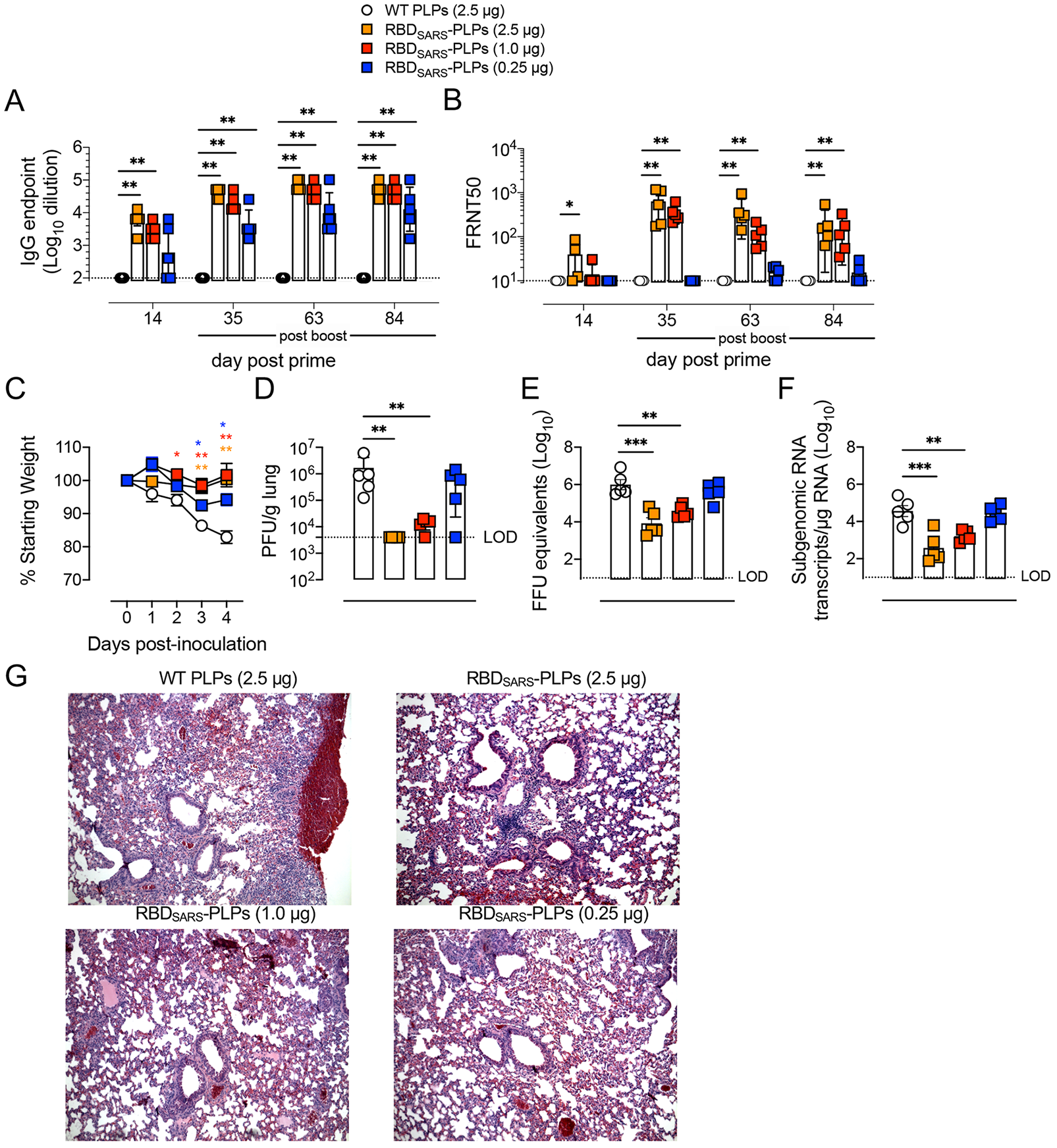
Immunization with RBD_SARS_-PLPs at low doses elicits potent and durable SARS-CoV-2 antibody responses and protective immunity. (**A-B**) WT BALB/c mice (n = 5/group) were immunized with 2.5 μg of WT PLPs (control) or variable doses (2.5, 1.0, or 0.25 μg) of 60% RBD_SARS_-PLPs by i.m. injection on days 0 and 21. Animals were bled on days 14, 35, 63, and 84. (**A**) RBD_SARS_-specific IgG endpoint titers were determined by ELISA. *P* values were determined by Mann-Whitney test. ^**^*P*<0.01. (**B**) SARS-CoV-2 neutralizing activity in serum samples was determined by FRNT. FRNT50 titer was calculated for each sample by nonlinear regression analysis. *P* values were determined by Mann-Whitney test. ^**^*P*<0.01. (**C**) At day 90 post-prime, mice were challenged i.n. with 10^4^ PFU of SARS-CoV-2 MA10 and monitored daily for changes in weight. *P* values were determined by two-way ANOVA with Tukey’s multiple comparisons test. ^*^*P*<0.05, ^**^*P*<0.01. (**D-G**) At 4 dpi, viral burden in the lung was quantified by plaque assay (**D**), RT-qPCR for viral genomic RNA (**E**), and RT-qPCR for N subgenomic mRNA (**F**), and histopathology in lung tissue sections was evaluated by hematoxylin and eosin staining (**G**). *P* values were determined by Mann-Whitney test. ^**^*P*<0.01, ^***^*P*<0.001.

We also evaluated serum from this dose de-escalation study for the capacity to neutralize SARS-CoV-2 infection (**Fig 6B and Fig S2**). Serum from mice immunized with WT PLPs did not display neutralizing activity (**Fig S2**). While day 14 post-prime serum from mice immunized with all doses of 60% RBD_SARS_-PLPs displayed little to no neutralizing activity, day 35 (14 days post-boost) serum from mice immunized with 2.5 or 1.0 μg of 60% RBD_SARS_-PLPs displayed potent neutralizing activity that was maintained up to 84 days post-prime (**Fig 6B**). Despite detectable levels of RBD_SARS_-specific IgG, immunization with 0.25 μg of 60% RBD_SARS_ PLPs did not result in neutralizing activity at any of the time points evaluated (**Fig 6B**).

Mice immunized with de-escalating doses of 60% RBD_SARS_ PLPs were challenged with 10^4^ PFU of SARS-CoV-2 MA10 at day 90 post-prime. Compared to control mice, mice immunized with all doses (including 0.25 μg) of 60% RBD_SARS_ PLPs were protected from weight loss associated with SARS-CoV-2 infection (**Fig 6C**). At 4 dpi, lungs were collected and assessed for viral burden. Immunization with 2.5 or 1.0 μg of 60% RBD_SARS_-PLPs resulted in potent protection against viral infection (**Fig 6D**) and reduced levels of viral genomic and subgenomic RNA (**Fig 6E,F**) in the lungs. Furthermore, histopathological analysis of lung tissues of mice immunized at these doses showed minimal perivascular and alveolar infiltrates at 4 dpi, whereas those of control mice were characterized by extensive inflammation (**Fig 6G**). Mice immunized with 0.25 μg of 60% RBD_SARS_ PLPs had levels of infectious virus and viral RNA in the lungs similar to those in control mice (**Fig 6D-F**), consistent with an inability to neutralize SARS-CoV-2 infection (**Fig 6B**). Additionally, these mice displayed more severe lung inflammation and injury versus those immunized with higher doses. Overall, the data indicate that neutralizing antibodies are essential for protection against SARS-CoV-2 infection and suggest that additional adaptive immune response elicited by vaccination with RBD_SARS_-PLPs may contribute to protection against severe disease characterized by more extensive weight loss, such as the generation of memory T cells.

### Vaccination with mosaic hCoV-PLPs protects from virulent SARS-CoV-2 and MERS-CoV challenge

As described above, PLPs can be decorated with multiple hCoV RBDs at varying surface densities to generate mosaic PLPs. Thus, we evaluated the immunogenicity and protective efficacy of particles simultaneously decorated with gpD-RBD_SARS_ and gpD-RBD_MERS_ in comparison with their mono-RBD decorated counterparts (**Fig 7**). Six-week-old BALB/c mice were immunized by i.m. injection with 7.5 μg of WT PLPs, 20% RBD_SARS_-PLPs, 20% RBD_MERS_-PLPs, or 20% hCoV-RBD-PLPs (note, 20% of each of gpD-RBD_SARS_ and gpD-RBD_MERS_). Mice received a booster dose of the same amount three weeks later. Serum samples were collected on days 14, 35 (14 days post-boost), and 96 post-prime, and IgG responses against purified SARS-CoV-2 and MERS-CoV RBD protein were evaluated by ELISA. At each time point evaluated, mice immunized with the bi-valent 20% hCoV-RBD-PLPs had similar levels of RBD_SARS_- and RBD_MERS_-specific IgG as mice immunized with the mono-valent 20% RBD_SARS_-PLPs or 20% RBD_MERS_-PLPs (**Fig 7A, 7B**). In addition, immunization with 20% hCoV-RBD-PLPs induced potent neutralizing antibody responses against both SARS-CoV-2 (**Fig 7C**) and MERS-CoV (**Fig 7D**). We note, however, that neutralizing titers against MERS-CoV were somewhat lower in mice immunized with the bi-valent vaccine compare with the mono-valent MERS vaccine (**Fig 7D**).

**Figure 7.**
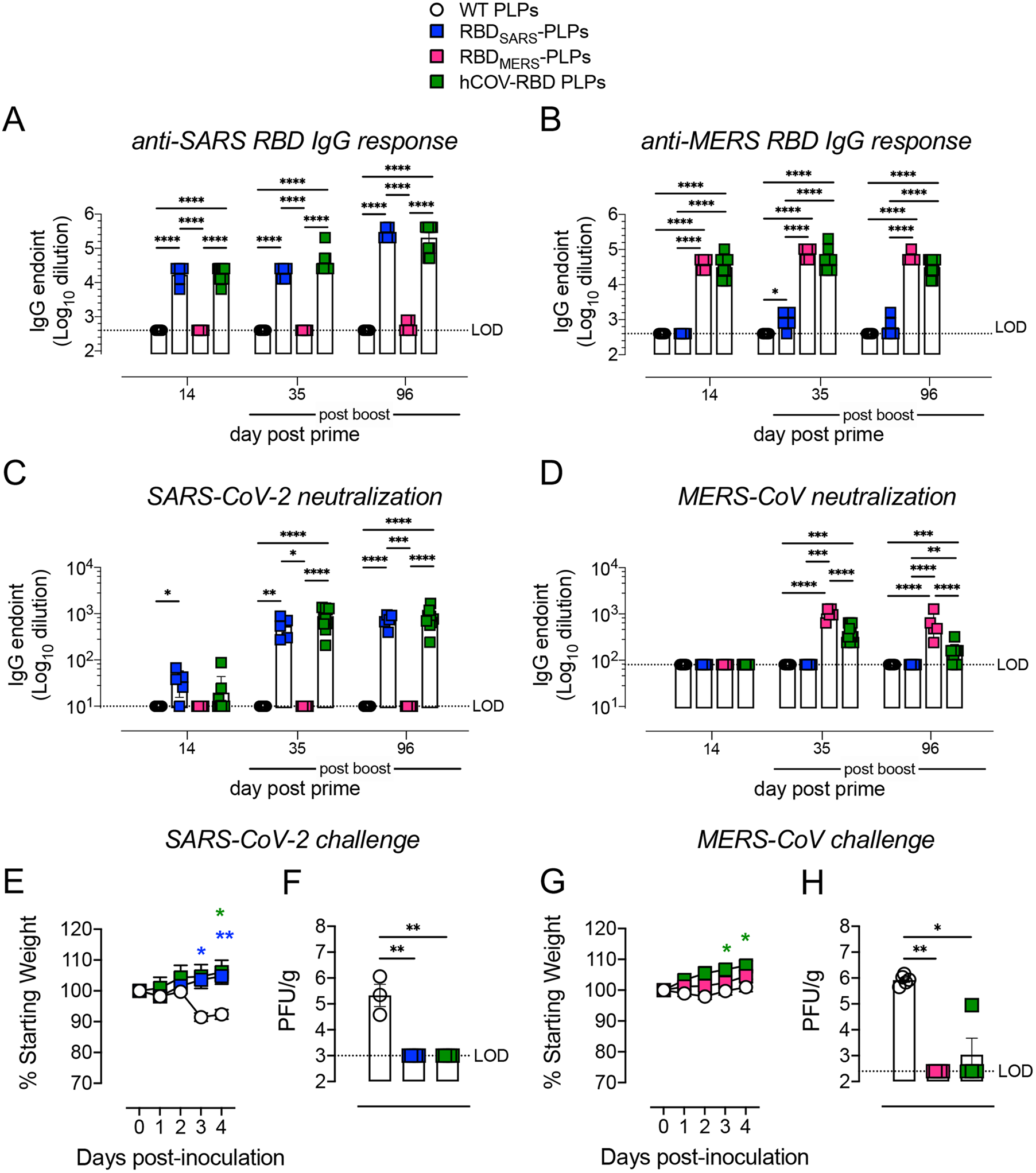
Immunization with mosaic hCoV-RBD-PLPs protects against both SARS-CoV-2 and MERS-CoV challenge. (**A-B**) WT BALB/c mice (n = 5-10 mice/group) were immunized with 7.5 μg of WT PLPs (control), 20% RBD_SARS_-PLPs, 20% RBD_MERS_-PLPs, or mosaic 20% hCoV-RBD-PLPs by intramuscular (i.m.) injection on days 0 and 21. At the times indicated, RBD_SARS_-specific (**A**) and RBD_MERS_-specific (**B**) IgG endpoint titers were determined by ELISA. Mann-Whitney test. ^*^*P*<0.05, ^***^*P*<0.001. (**C**) SARS-CoV-2 neutralizing activity was determined by a focus reduction neutralization test (FRNT). The dilution of serum that inhibited 50% of infectivity (FRNT50 titer) was calculated. Mann-Whitney test. ^*^*P*<0.05, ^**^*P*<0.01, ^***^*P*<0.001, ^****^*P*<0.0001. (**D**) MERS-CoV neutralizing activity was determined by a plaque reduction neutralization test (PRNT). The dilution of serum that inhibited 50% of infectivity (PRNT50 titer) was calculated. Mann-Whitney test. ^**^*P*<0.01, ^***^*P*<0.001, ^****^*P*<0.0001. (**E-H**) At day 102 post-prime, mice were challenged intranasally (i.n.) with 10^4^ PFU of SARS-CoV-2 MA10 (**E-F**) or 10^5^ PFU MERS-CoV (**G-H**). (**E and G**) Mice were monitored daily for weight changes. Two-way ANOVA with Tukey’s multiple comparisons test. ^**^*P*<0.01, ^**^*P*<0.01. (**F and H**) At 4 dpi, the burden of infectious SARS-CoV-2 (**F**) and MERS-CoV (**H**) in the lung was quantified by plaque assay. Mann-Whitney test. ^*^*P*<0.05, ^**^*P*<0.01.

Next, these mice were challenged with 10^4^ PFU of SARS-CoV-2 MA10 or 10^5^ PFU MERS-CoV at day 102 post-prime. Five days prior to MERS-CoV challenge, mice were i.n. inoculated with Ad-hDPP4 to deliver the MERS-CoV cell entry receptor to lung cells (*56, 57*). Compared with control mice, mice immunized with either RBD_SARS_-PLPs or hCoV-PLPs were protected from weight loss (**Fig 7E**) and lung viral burden (**Fig 7F**) associated with SARS-CoV-2 infection. Similarly, mice immunized with either RBD_MERS_-PLPs or hCoV-PLPs were protected from weight loss (**Fig 7G**) and lung viral burden (**Fig 7H**) associated with MERS-CoV infection.

## DISCUSSION

Numerous vaccine strategies and platforms have been developed for current and emerging infectious diseases, and several studies support the use of CoV RBDs as vaccine immunogens. For instance, the SARS-CoV-2 RBD is immunodominant, contains multiple antigenic sites, and is the overwhelming target of neutralizing activity in COVID-19 convalescent plasma (*58*). In addition, passive administration of monoclonal antibodies that target the RBD of SARS-CoV-2 and other pathogenic CoVs protects rodents, ferrets, and non-human primates from severe CoV infection (*14, 59-68*), indicating that the generation of RBD-specific antibodies is an advantageous therapeutic strategy. Moreover, the RBD is a target of cross-neutralizing antibodies (*58, 69*), and immunization with SARS-CoV-2 RBD functionalized nanoparticles can elicit cross-neutralizing antibodies against a number of SARS-related CoVs (*70*). Nevertheless and despite significant advancements in diagnostic and therapeutic countermeasures, continued vaccine development efforts are warranted, especially for viruses that pose a threat to global public health or to provide broad protection against multiple viruses.

In this study, we designed and constructed lambda PLPs decorated with the RBDs from extant highly pathogenic human CoVs (i.e., SARS-CoV-2, MERS-CoV) as a tunable, multivalent vaccine platform. The platform allows for symmetrical, high-density display of multiple target antigens, thereby enhancing the stimulation of host immune responses. Properties of decorated PLPs present several advantages for use as an effective vaccine delivery vehicle, including (i) a particle size less than 200 nm, enabling entry into the lymphatic system (*71*) where adaptive immune responses are initiated; (ii) additional physiochemical properties such as surface charge, exposure of functional group, and hydrophobicity are easily manipulated by both genetic and chemical approaches; and (iii) simultaneous delivery of foreign molecules can be achieved on a single PLP. Further, we found no adverse effects due to immunization with phage lambda particles by intramuscular routes in mice even after repeated doses or after challenge (i.e., no evidence of vaccine-enhanced disease), indicating a favorable safety profile for future *in vivo* testing.

We demonstrate that mono and mosaic hCoV-RBD decorated particles induce RBD-specific binding and virus neutralizing antibodies after one and two doses. Notably, these responses remained highly elevated up to 6 months post-prime, indicating that the vaccine platform can stimulate robust and durable immune responses. Additionally, immunized mice were protected against virulent virus challenge (up to day 184 post-prime for SARS-CoV-2 and day 102 for MERS-CoV) as indicated by significantly reduced virus-induced weight loss, a marked reduction in lung-associated inflammation and injury, and lower titers of infectious virus in the lungs paired with diminished levels of viral genomic and subgenomic RNA. Consistent with published studies (*70*), PLPs decorated with the SARS-CoV-2 RBD alone do not elicit IgG that binds the MERS-CoV RBD or antibodies that neutralize MERS-CoV infection. In contrast, those simultaneously decorated with the SARS-CoV-2 and MERS-CoV RBD proteins induce strong immune responses against both pathogenic viruses. This supports that co-displaying multiple antigens on the same particle is an effective means of broadening immune responses elicited by our vaccine platform, and that the presence of multiple antigens do not interfere with each other.

A number of nanoparticles and antigen coupling strategies have been evaluated for vaccine design (*72*). These include nanoparticles based on the SpyCatcher-SpyTag system which can be used to conjugate antigens to self-assembling protein nanoparticles (*73-75*), the Novavax vaccine (*76*), and other phage systems (*77*). Most often, these immunization studies involve the use of large amounts of antigen (e.g., 50-100 μg/dose), up to 4 immunizations, formulation with an adjuvant (e.g., AddaVax), and vaccine administration routes not amenable to humans (e.g., intraperitoneal) (*75, 78*). Our dose response studies show that immunization with as little as one microgram of 60% RBD_SARS_-PLPs elicited robust neutralizing antibody responses and conferred protection against SARS-CoV-2 challenge. These data suggest that PLP-based vaccines facilitate dose sparing immunization practices which, from both manufacturing and distribution perspectives, enhances their utility as a vaccine platform. Further studies will be necessary to test protective efficacy following immunization in a dose-dependent manner or through alternative routes.

While we did not define the epitopes targeted, the anti-SARS-CoV-2 monoclonal antibody used in the enzyme immunoassays binds to a conserved epitope in the RBD that is only accessible when the S protein is in conformation competent to bind the hACE2 receptor (*79, 80*). Additionally, RBD-specific IgG subclass responses were assessed; however, further characterization of the potential role of these in vaccine-mediated protection was not determined. Considering the implications these may have to the development of safe vaccines (e.g., clinical syndromes associated with vaccine-enhanced disease) (*81*), additional studies are needed to examine the possible impacts to cellular effector function and evaluate mucosal and cellular immunity in addition to immunity against variants of SARS-CoV-2 and MERS-CoV.

In summary, the versatility and robustness of the lambda system presented provides a promising vaccine candidate. Immunization with hCoV-RBD decorated PLPs induces antibodies with neutralizing activity, results in a durable immune response, and provides effective protection against virulent virus challenge. Continued efforts into the optimization of this platform could expand the diversity of protein-based vaccine technologies available and aid in the prevention of infectious diseases deleterious to global health.

## METHODS

### Purification of lambda PLPs

Lambda PLPs were expressed in *E. coli* BL21(DE3)[pNu3_E] cells and purified, as described (*46*). Briefly, self-assembled PLPs were extracted from cell lysates in 20 mM Tris [pH 8.0, 4°C] buffer containing 0.4 mg/mL lysozyme and 0.04 mg/mL DNase. Following rate zonal centrifugation (10-40% sucrose density gradient), PLPs were collected, concentrated, and exchanged into 20 mM Tris [pH 8.0, 4°C] buffer containing 15 mM MgCl_2_, 1 mM EDTA, and 7 mM β-ME using centrifugal filter units (100k MWCO; Millipore Amicon Ultra). Proteins were fractionated by anion exchange chromatography employing three 5 mL HiTrap Q HP columns connected in tandem and developed with a 30-column volume linear gradient to 1 M NaCl. The eluate was analyzed by SDS-PAGE, and PLP-containing fractions were pooled and exchanged into 50 mM HEPES [pH 7.4] buffer containing 100 mM NaCl and 10 mM MgCl_2_ prior to storage at 4 °C.

### Purification of lambda decoration proteins

WT and gpD(S42C) decoration proteins were expressed in *E. coli* BL21(DE3)[pD] and BL21(DE3)[pDS542C] cells, respectively, and purified, as described (*46*). Briefly, cell lysate supernatants were dialyzed overnight against 20 mM Tris [pH8.0, 4 °C] buffer containing 20 mM NaCl and 0.1 mM EDTA. Proteins were fractionated employing three 5 mL HiTrap Q HP columns connected in tandem and developed with a 30-column volume linear gradient to 1 M NaCl. Fractions containing decoration protein were pooled, exchanged into 50 mM NaOAc [pH 4.8] buffer using centrifugal filter units (3k MWCO; Millipore Amicon Ultra), and loaded onto three 5 mL HiTrap SP columns connected in tandem. Bound proteins were eluted with a 30-column volume linear gradient to 0.5 M NaCl, and fractions containing decoration protein were pooled and dialyzed overnight against 20 mM Tris [pH 8.0, 4 °C] buffer containing 20 mM NaCl and 0.1 mM EDTA for storage at 4°C. For long-term storage at -80°C, aliquots of purified protein were supplemented with 20% glycerol.

### Purification of recombinant hCoV RBD proteins

A pCAGGS expression vector encoding the SARS-CoV-2 spike RBD was obtained from Dr. Florian Krammer, Icahn School of Medicine at Mount Sinai (*82*). This construct contains sequences encoding the native signal peptide (residues 1–14) followed by residues 319–541 from the SARS-CoV-2 Wuhan-Hu-1 spike protein (GenBank MN908947.3) and a hexa-histidine tag at the C-terminus for purification. A pTwist-CMV expression vector (Twist Biosciences Technology) encoding the MERS-CoV spike RBD was kindly provided by Dr. Peter S. Kim, Stanford University (*83*). This construct contains sequences encoding an N-terminal human IL-2 signal peptide following by residues 367-588 from the MERS-CoV EMC/2012 spike RBD (GenBank JX86905.2) and dual C-terminal tags (octa-histidine, AviTag) for purification. Recombinant protein expression and purification was performed by the University of Colorado Cancer Center Cell Technologies Shared Resource. Due to the involvement of glycans in the epitopes of some neutralizing antibodies (*62, 84*), RBDs were expressed in Expi293F cells to retain normal glycosylation and antigenic properties. Expi293F cells were transfected with plasmid DNA using ExpiFectamine transfection reagent (ThermoFisher Scientific). At 72 h post-transfection, cell culture supernatants were centrifuged (4,000 x g) for 20 min and filtered (0.2 micron). Recombinant proteins were purified by Ni-NTA column chromatography using an ATKA purification system. Protein was eluted with 500 mM imidazole and concentrated before assessing protein purity by SDS-PAGE and Coomassie staining.

### Crosslinking of lambda decoration protein (gpD(S42C)) to hCoV RBDs

SARS-CoV-2 or MERS-CoV RBD protein was exchanged into 0.01 M PBS [pH 7.2] buffer using centrifugal filter units (10k MWCO; Millipore Amicon Ultra). A 5-fold molar excess of SM(PEG)_24_ (PEGylated, long-chain sulfosuccinimidyl 4-(N-maleimidomethyl)cyclohexane-1-carboxylate crosslinker; ThermoFisher Scientific) was added, and the mixture was incubated (30 min, 25°C) for modification of solvent accessible primary amines. Reduced gpD(S42C) was added to the mixture (1:1, molar equivalent) and incubated (1 h, 25°C) to yield the crosslinked products (gpD-RBD_SARS_, gpD-RBD_MERS_). The reaction was quenched with the addition of 0.2% β-ME (30 min, 25°C), and crosslinked proteins were purified by size-exclusion chromatography employing a 24 mL Superose 6 Increase 10/300 GL column equilibrated and developed with 40 mM HEPES [pH 7.4] buffer containing 150 mM NaCl, 200 mM arginine, 0.1 mM EDTA, and 2 mM β-ME. Fractions containing cross-linked proteins were identified by SDS-PAGE and were pooled. The purified proteins were and exchanged into 20 mM Tris [pH 8.0, 4°C] buffer containing 20 mM NaCl and 0.1 mM EDTA for storage at 4°C. Protein concentration was quantified spectroscopically.

### *In vitro* PLP expansion and decoration

PLPs were expanded and subsequently decorated *in vitro*, as described (*46*). Briefly, purified PLPs were expanded in 10 mM HEPES [pH 7.4] buffer containing 2.5 M urea for 30 min on ice and then exchanged into 10 mM HEPES [pH 7.4] buffer containing 200 mM urea using centrifugal filter units (100k MWCO; Millipore Amicon Ultra). Expanded shells (30 nM) were decorated in a stepwise fashion with modified and WT decoration protein at 25°C in 10 mM HEPES [pH 7.4] buffer containing 50 mM urea, 10 mM arginine, and 0.1% Tween 20. Proteins were added in the following order: (1) gpD-RBD_SARS_ (0.69-8.33 μM final concentration, 30 min incubation); (2) gpD-RBD_MERS_ (1.39-8.33 μM, 30 min incubation); (3) wildtype gpD (5.55-13.19 μM, 60 min incubation). Decorated PLPs were purified by size-exclusion chromatography employing a 24 mL Superose 6 Increase 10/300 GL column equilibrated and developed with 100 mM HEPES [pH 6.6] buffer containing 200 mM NaCl, 200 mM arginine, 1 mM EDTA, and 2 mM β-ME at a flow rate of 0.3 mL/min. Fractions containing decorated PLPs were pooled and exchanged into 50 mM HEPES [pH 7.4] buffer containing 100 mM NaCl and 10 mM MgCl_2_ for storage at 4°C.

### Transmission Electron Microscopy

Carbon-coated copper grids (400 mesh; CF400-CU) were glow-discharged using a Pelco EasiGlo system (Ted Pella, Inc.; Redding, CA, USA) with a plasma current of 15 mA, negative glow discharge head polarity, glow discharge duration of one minute, and held under vacuum for 15 seconds. PLP preparations were diluted to 10 nM using water (sterile, double distilled) and spotted twice onto grids. Following sample adsorption (20 s), excess liquid was wicked off using a Whatman #1 filter paper. Grids were washed using water, and excess liquid was wicked off. Samples were negatively stained (20-25 s) with filtered 2% (w/v) methylamine tungstate [pH 6.7] and 50 μg/mL bacitracin (0.2 μm Nucleopore polycarbonate syringe filters). Excess stain was wicked off, and grids were allowed to air-dry for at least one hour before transferring to a grid storage box. Samples were maintained covered throughout this procedure and stored at room temperature until imaged. Images were acquired on a FEI Tecnai G2 transmission electron microscope at an accelerating voltage of 80 kV and equipped with a 2k × 2k CCD camera. Images were processed in Fiji (*85*) and measurements were based on 100 particles with values reported as mean ± standard deviation.

### Particle size and charge measurements

Decorated particles were diluted to 2 nM (55-700 μL) using 10 mM HEPES [pH 7.4] for dynamic light scattering (DLS) and electrophoretic light scattering (ELS) analyses. Particle size and charge measurements were acquired on a Malvern Panalytical Zetasizer Nano ZS (He-Ne laser 633 nm light source; 5 mW maximum power). Size information obtained from the correlation function was based on hydrodynamic diameter (weighted mean reported as Z-average, Z-Ave) and particle size distributions (intensity, volume PSD) with polydispersity index (PDI) values provided to serve as an estimate of the width of the distributions. Overall surface charge was reported as zeta potential [mV]. Successive sample measurements were done in quadruplicate with values reported as mean ± standard deviation.

### Virus and cells

Vero E6 (CRL-1586, American Type Culture Collection (ATCC)) were cultured at 37°C in Dulbecco’s Modified Eagle medium (DMEM, HyClone 11965-084) supplemented with 10% fetal bovine serum (FBS), 10 mM HEPES [pH 7.3], 1 mM sodium pyruvate, 1X non-essential amino acids, and 100 U/ml of penicillin–streptomycin. SARS-CoV-2 strain 2019 n-CoV/USA_WA1/2020 was obtained from BEI Resources. The virus was passaged once in Vero E6 cells and titrated by focus formation assay (FFA) on Vero E6 cells. The mouse-adapted SARS-CoV-2 strain MA10 (kindly provided by R. Baric, University of North Carolina at Chapel Hill) elicits disease signs in laboratory mice similar to severe COVID-19 (*55*) was used for all challenge studies. All work with infectious SARS-CoV-2 and MERS-CoV was performed in Institutional Biosafety Committee approved BSL3 and A-BSL3 facilities at the University of Colorado School of Medicine and University of Maryland School of Medicine using positive pressure air respirators and other personal protective equipment.

### Mouse experiments

Animal studies were carried out in accordance with the recommendations in the Guide for the Care and Use of Laboratory Animals of the National Institutes of Health. The protocols were approved by the Institutional Animal Care and Use Committees at the University of Colorado School of Medicine (Assurance Number A3269-01) and the University of Maryland School of Medicine (Assurance number D16-00125 (A3200-01)). Female BALB/c mice were purchased from The Jackson Laboratory. 6-8-week-old mice were immunized with WT PLPs (control), RBD_SARS_-PLPs, RBD_MERS_-PLPs, or hCoV-RBD PLPs preparations in 50 μl PBS via i.m. injection in the hind leg. Mice were boosted three weeks after the primary immunization using the same route. For virulent virus challenge, mice were anaesthetized by intraperitoneal injection with 50 μL of a mix of xylazine (0.38 mg/mouse) and ketamine hydrochloride (1.3 mg/mouse) diluted in PBS. Immunized BALB/c mice were inoculated intranasally (i.n.) with 10^4^ PFU of SARS-CoV-2 MA10. For MERS-CoV challenge, immunized mice were first inoculated i.n. with 2.5 × 10^8^ PFU of a replication incompetent adenovirus vector to deliver human DPP4 (Ad5-hDPP4) into the lungs of mice (*57*). Five days later, mice were inoculated i.n. with 10^5^ PFU of MERS-CoV (Jordan strain). All mice were weighed and monitored for additional disease signs daily. At 4 days post-inoculation (dpi), animals were euthanized by bilateral thoracotomy and tissues were collected for virological, immunological, and pathological analyses.

### ELISA

SARS-CoV-2 and MERS-CoV RBD-specific antibody responses in mouse and human sera were measured by ELISA. Immunol 4HBX plates were coated with 0.2 μg of recombinant SARS-CoV-2 RBD protein (Wuhan-Hu-1, GenPept QHD43416) or MERS-CoV RBD protein overnight at 4°C. Coating antigen was removed, and plates were blocked with 200 μl of 3% non-fat milk in PBS/0.1% Tween-20 (PBS-T) for one hour at room temperature. After blocking, wells were washed once with PBS-T and then probed with serum samples diluted in PBS-T supplemented with 1% non-fat milk for 1.5 h. Samples were removed, and wells were washed three times with PBS-T and probed with secondary antibodies diluted at 1:4000 in PBS-T; goat anti-mouse IgG-HRP (Southern Biotech, 1030-05), goat anti-human IgG Fc-HRP (Southern Biotech, 2014-05), goat anti-mouse IgG1-BIOT (Southern Biotech, 1071-08), goat anti-mouse IgG_2a_ Human ads-BIOT (Southern Biotech, 1080-08), goat anti-mouse IgG_2b_-BIOT (Southern Biotech, 1091-08), goat anti-mouse IgG_3_ human ads-BIOT (Southern Biotech, 1100-08). Biotinylated antibodies were subsequently probed with streptavidin-HRP (Southern Biotech, 7100-05). Detection antibody was removed, and wells were washed three times with PBS-T. Plates were developed using 3,3’,5,5’-tetramethylbenzidine (TMB; Sigma, T0440), followed by the addition of 0.3 M H_2_SO_4_. Plate absorbance was read at 450 nm on a Tecan Infinite M plex plate reader. Endpoint titers are reported as the reciprocal of the final dilution, in which absorbance at 450 nm was above the blank average plus three times the standard deviation of all blanks in the assay. The immunoreactivity of RBD_SARS_-PLPs, RBD_MERS_-PLPs, and hCoV-RBD PLPs was measured in a similar manner. In this case, plates were coated with 0.2 μg of WT PLPs or RBD_SARS_-PLPs, RBD_MERS_-PLPs, and hCoV-RBD PLPs overnight at 4°C. Following blocking and washing, wells were incubated for 1.5 h at room temperature with either chimeric human anti-SARS-CoV spike antibody clone CR3022 (Absolute Antibody, Ab01680) or mouse anti-MERS-CoV spike antibody clone D12 (Absolute Antibody, Ab00696) prepared in a 2-fold dilution series in PBS-T with a starting dilution of 1:200 and signal was developed as described above.

### SARS-CoV-2 focus reduction neutralization test (FRNT)

Vero E6 cells were seeded in 96-well plates at 10^4^ cells/well. The next day serum samples were heat-inactivated at 56°C for 30 minutes and then serially diluted (2-fold, starting at 1:10) in DMEM (HyClone, 11965-084) plus 1% FBS in 96-well plates. Approximately 100 focus-forming units (FFU) of SARS-CoV-2 USA-WA1/2020 were added to each well and the serum plus virus mixture was incubated for 1 h at 37°C. Following co-incubation, medium was removed from cells and the serum sample plus virus mixture was added to the cells for 1 h 37°C. Samples were then removed and cells overlaid with 1% methylcellulose (MilliporeSigma, M0512) in minimum essential media (MEM) (Gibco, 12000-063) plus 2% FBS and incubated for 24 hours at 37°C. Cells were fixed with 1% paraformaldehyde (PFA; Acros Organics, 416780030) and probed with 1 μg/mL of chimeric human anti-SARS-CoV spike antibody (CR3022, Absolute Antibody, Ab01680) in Perm Wash (1X PBS/0.1% saponin/0.1% BSA) for 2 h at room temperature. After three washes with PBS-T, cells were incubated with goat anti-human IgG Fc-HRP (Southern Biotech, 2014-05) diluted at 1:1000 in Perm Wash for 1.5 h at room temperature. SARS-CoV-2-positive foci were visualized with TrueBlue substrate (SeraCare, 5510-0030) and counted using a CTL Biospot analyzer and Biospot software (Cellular Technology Ltd, Shaker Heights, OH). The FRNT_50_ titers were calculated relative to a virus only control (no serum) set at 100%, using GraphPad Prism 9.1.2 (La Jolla, CA) default nonlinear curve fit constrained between 0 and 100%.

### MERS-CoV neutralization assay

To determine the inhibitory activity of mouse sera against MERS-CoV, 3,950 TCID_50_/ml of MERS-COV-Jordan was incubated with diluted sera for 30 minutes at room temperature and the inhibitory capacity of each serum dilution was assessed by TCID_50_ assay as previously described (*56, 86*).

### Quantification of SARS-CoV-2 genomic and subgenomic RNA

To quantify viral genomic and subgenomic RNA, lung tissue of mice challenged with SARS-CoV-2 was homogenized in TRIzol reagent (Life Technologies, 15596018) and total RNA was isolated using a PureLink RNA Mini kit (Life Technologies, 12183025). Single-stranded cDNA was generated using random primers and SuperScript IV reverse transcriptase (Life Technologies, 18091050). SARS-CoV-2 genomic or subgenomic RNA copies were measured by qPCR using the primer and probe combinations listed in **Table S1** (Integrated DNA Technologies). To quantify SARS-CoV-2 genomic RNA, we extrapolated viral RNA levels from a standard curve generated from known FFU of SARS-CoV-2 from which RNA was isolated and cDNA generated as previously described (*87*). To quantify SARS-CoV-2 subgenomic RNA, we extrapolated viral RNA levels from a standard curve using defined concentrations of a plasmid containing an amplified SARS-CoV-2 subgenomic fragment (pCR-sgN TOPO). Briefly, a subgenomic RNA fragment was amplified by RT-PCR using RNA isolated from SARS-CoV-2-infected Vero E6 cells. The 125 bp amplicon is composed of the joining region between the 3’ UTR and the N gene, with inclusion of a transcription-regulatory sequence (TRS) of the SARS-CoV-2 Wuhan-Hu-1 (NC_045512.2). This amplicon was cloned into the pCR4 Blunt TOPO vector (Invitrogen, K2875J10), sequence confirmed, and used in a dilution series of defined gene copies. All qPCR reactions were prepared with Taqman Universal MasterMix II (Applied Biosystems, 4440038) and were analyzed with Applied Biosystems QuantStudio ViiA 7 analyzer.

### Plaque assay

Vero E6 cells were seeded in 12-well plates one day prior to virus inoculation. Lung homogenates were serially diluted in DMEM supplemented with 2% FBS, HEPES, and penicillin-streptomycin and incubated on cells for one hour at 37°C. Afterwards, cells were overlaid with 1% (w/v) methylcellulose in MEM plus 2% FBS and incubated at 37°C. At 3 dpi, overlays were removed, and plates were fixed with 4% PFA for 20 minutes at room temperature. After removal of PFA, plates were stained with 0.05% (w/v) crystal violet in 20% methanol (10-20 min). Crystal violet was removed, and plates were rinsed with water or PBS. Plaques were counted manually to determine infectious virus titer.

### Histopathology

Mice were euthanized at 4 days following SARS-CoV-2 MA10 or MERS-CoV challenge. The lungs were removed and fixed with 10% formalin. 5-micron sections were stained with H&E for histological examination. Slides were examined in a blinded fashion for total inflammation, periarteriolar, and peribronchiolar inflammation and epithelial cell denuding.

### Statistical Analysis

Statistical significance was assigned when *P value*s were < 0.05 using Prism Version 9.1.2 (GraphPad). Tests, number of animals (n), median values, and statistical comparison groups are indicated in the Figure Legends.

## ACKNOWLEDGEMENTS

The authors gratefully acknowledge Florian Krammer and Peter S. Kim for providing the expression vectors for the SARS-CoV-2 spike RBD and MERS-CoV spike RBD proteins, respectively. We thank Ralph S. Baric for providing the mouse-adapted SARS-CoV-2 strain MA10

The research reported herein was supported by the NSF #GRFP1553798 (A.C.) and #MCB2016019 (C.E.C.) as well as funds from the University of Colorado School of Medicine.

## AUTHOR CONTRIBUTIONS

B.J.D., A.C., M.B.F., C.E.C., and T.E.M. designed the experiments. B.J.D., A.C., S.W., and R.J. performed the experiments. B.J.D., A.C., S.W., R.J., M.B.F., C.E.C., and T.E.M. performed data analysis. B.J.D., A.C., C.E.C., and T.E.M. wrote the initial draft of the manuscript. All authors provided comments and edits to the final version.

## FIGURE LEGENDS

**TABLE S1:**
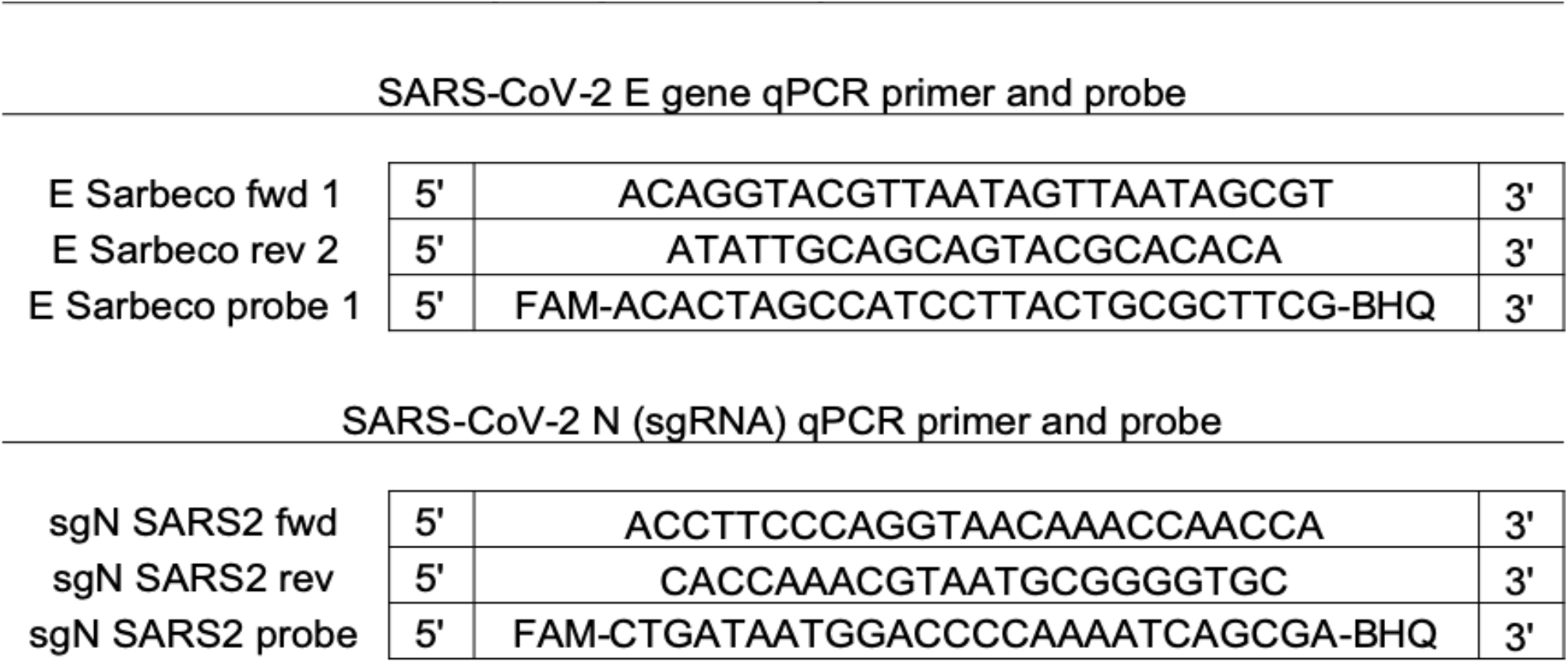
SARS-CoV-2 qPCR Primer and Probe combinations.

## REFERENCES

1. N. Zhu et al., A Novel Coronavirus from Patients with Pneumonia in China, 2019. N Engl J Med 382, 727–733 (2020).

2. F. Wu et al., A new coronavirus associated with human respiratory disease in China. Nature 579, 265–269 (2020).

3. V. Coronaviridae Study Group of the International Committee on Taxonomy of, The species Severe acute respiratory syndrome-related coronavirus: classifying 2019-nCoV and naming it SARS-CoV-2. Nat Microbiol 5, 536–544 (2020).

4. K. R. Stenmark et al., Mechanisms of SARS-CoV-2-induced lung vascular disease: potential role of complement. Pulm Circ 11, 20458940211015799 (2021).

5. N. Stefan, A. L. Birkenfeld, M. B. Schulze, Global pandemics interconnected - obesity, impaired metabolic health and COVID-19. Nat Rev Endocrinol, (2021).

6. Z. A. Memish, S. Perlman, M. D. Van Kerkhove, A. Zumla, Middle East respiratory syndrome. Lancet 395, 1063–1077 (2020).

7. A. M. Zaki, S. van Boheemen, T. M. Bestebroer, A. D. Osterhaus, R. A. Fouchier, Isolation of a novel coronavirus from a man with pneumonia in Saudi Arabia. N Engl J Med 367, 1814–1820 (2012).

8. E. Petersen, D. S. Hui, S. Perlman, A. Zumla, Middle East Respiratory Syndrome - advancing the public health and research agenda on MERS - lessons from the South Korea outbreak. Int J Infect Dis 36, 54–55 (2015).

9. M. S. Majumder, C. Rivers, E. Lofgren, D. Fisman, Estimation of MERS-Coronavirus Reproductive Number and Case Fatality Rate for the Spring 2014 Saudi Arabia Outbreak: Insights from Publicly Available Data. PLoS Curr 6, (2014).

10. P. V’Kovski, A. Kratzel, S. Steiner, H. Stalder, V. Thiel, Coronavirus biology and replication: implications for SARS-CoV-2. Nat Rev Microbiol 19, 155–170 (2021).

11. F. Li, Structure, Function, and Evolution of Coronavirus Spike Proteins. Annu Rev Virol 3, 237–261 (2016).

12. M. Hoffmann et al., SARS-CoV-2 Cell Entry Depends on ACE2 and TMPRSS2 and Is Blocked by a Clinically Proven Protease Inhibitor. Cell 181, 271–280 e278 (2020).

13. B. Ju et al., Human neutralizing antibodies elicited by SARS-CoV-2 infection. Nature 584, 115–119 (2020).

14. S. J. Zost et al., Potently neutralizing and protective human antibodies against SARS-CoV-2. Nature 584, 443–449 (2020).

15. L. Premkumar et al., The receptor binding domain of the viral spike protein is an immunodominant and highly specific target of antibodies in SARS-CoV-2 patients. Sci Immunol 5, (2020).

16. L. Liu et al., Potent neutralizing antibodies against multiple epitopes on SARS-CoV-2 spike. Nature 584, 450–456 (2020).

17. J. Hansen et al., Studies in humanized mice and convalescent humans yield a SARS-CoV-2 antibody cocktail. Science 369, 1010–1014 (2020).

18. Y. Li et al., A humanized neutralizing antibody against MERS-CoV targeting the receptor-binding domain of the spike protein. Cell Res 25, 1237–1249 (2015).

19. X. C. Tang et al., Identification of human neutralizing antibodies against MERS-CoV and their role in virus adaptive evolution. Proc Natl Acad Sci U S A 111, E2018–2026 (2014).

20. L. Jiang et al., Potent neutralization of MERS-CoV by human neutralizing monoclonal antibodies to the viral spike glycoprotein. Sci Transl Med 6, 234ra259 (2014).

21. K. E. Pascal et al., Pre- and postexposure efficacy of fully human antibodies against Spike protein in a novel humanized mouse model of MERS-CoV infection. Proc Natl Acad Sci U S A 112, 8738–8743 (2015).

22. L. Du et al., Identification of a receptor-binding domain in the S protein of the novel human coronavirus Middle East respiratory syndrome coronavirus as an essential target for vaccine development. J Virol 87, 9939–9942 (2013).

23. F. Krammer, SARS-CoV-2 vaccines in development. Nature 586, 516–527 (2020).

24. J. P. Moore, P. J. Klasse, COVID-19 Vaccines: “Warp Speed” Needs Mind Melds, Not Warped Minds. J Virol 94, (2020).

25. F. P. Polack et al., Safety and Efficacy of the BNT162b2 mRNA Covid-19 Vaccine. N Engl J Med 383, 2603–2615 (2020).

26. L. R. Baden et al., Efficacy and Safety of the mRNA-1273 SARS-CoV-2 Vaccine. N Engl J Med 384, 403–416 (2021).

27. J. Sadoff et al., Safety and Efficacy of Single-Dose Ad26.COV2.S Vaccine against Covid-19. N Engl J Med 384, 2187–2201 (2021).

28. J. Wise, Covid-19: European countries suspend use of Oxford-AstraZeneca vaccine after reports of blood clots. BMJ 372, 699 (2021).

29. H. Boytchev, Covid-19: Germany struggles with slow uptake of Oxford AstraZeneca vaccine. BMJ 372, 619 (2021).

30. E. J. Kremer, Pros and Cons of Adenovirus-Based SARS-CoV-2 Vaccines. Mol Ther 28, 2303–2304 (2020).

31. K. J. Koudelka, A. S. Pitek, M. Manchester, N. F. Steinmetz, Virus-Based Nanoparticles as Versatile Nanomachines. Annu Rev Virol 2, 379–401 (2015).

32. M. Karimi et al., Bacteriophages and phage-inspired nanocarriers for targeted delivery of therapeutic cargos. Adv Drug Deliv Rev 106, 45–62 (2016).

33. P. Tao, J. Zhu, M. Mahalingam, H. Batra, V. B. Rao, Bacteriophage T4 nanoparticles for vaccine delivery against infectious diseases. Adv Drug Deliv Rev 145, 57–72 (2019).

34. K. A. Kelly, P. Waterman, R. Weissleder, In vivo imaging of molecularly targeted phage. Neoplasia 8, 1011–1018 (2006).

35. Z. M. Carrico et al., N-Terminal labeling of filamentous phage to create cancer marker imaging agents. ACS Nano 6, 6675–6680 (2012).

36. J. L. Liu, K. L. Robertson, Engineered bacteriophage T4 nanoparticles for cellular imaging. Methods Mol Biol 1108, 187–199 (2014).

37. T. Yata et al., Hybrid Nanomaterial Complexes for Advanced Phage-guided Gene Delivery. Mol Ther Nucleic Acids 3, e185 (2014).

38. M. E. Farkas et al., PET Imaging and biodistribution of chemically modified bacteriophage MS2. Mol Pharm 10, 69–76 (2013).

39. J. R. Chang et al., Phage lambda capsids as tunable display nanoparticles. Biomacromolecules 15, 4410–4419 (2014).

40. N. E. van Houten, K. A. Henry, G. P. Smith, J. K. Scott, Engineering filamentous phage carriers to improve focusing of antibody responses against peptides. Vaccine 28, 2174–2185 (2010).

41. B. Schwarz et al., Symmetry Controlled, Genetic Presentation of Bioactive Proteins on the P22 Virus-like Particle Using an External Decoration Protein. ACS Nano 9, 9134–9147 (2015).

42. P. Tao et al., In vitro and in vivo delivery of genes and proteins using the bacteriophage T4 DNA packaging machine. Proc Natl Acad Sci U S A 110, 5846–5851 (2013).

43. K. D. Brune et al., Plug-and-Display: decoration of Virus-Like Particles via isopeptide bonds for modular immunization. Sci Rep 6, 19234 (2016).

44. N. Stephanopoulos, G. J. Tong, S. C. Hsiao, M. B. Francis, Dual-surface modified virus capsids for targeted delivery of photodynamic agents to cancer cells. ACS Nano 4, 6014–6020 (2010).

45. A. Razazan et al., Lambda bacteriophage nanoparticles displaying GP2, a HER2/neu derived peptide, induce prophylactic and therapeutic activities against TUBO tumor model in mice. Sci Rep 9, 2221 (2019).

46. A. Catala et al., Targeted Intracellular Delivery of Trastuzumab Using Designer Phage Lambda Nanoparticles Alters Cellular Programs in Human Breast Cancer Cells. ACS Nano, (2021).

47. C. E. Catalano, Bacteriophage lambda: The path from biology to theranostic agent. Wiley Interdiscip Rev Nanomed Nanobiotechnol 10, e1517 (2018).

48. G. C. Lander et al., Bacteriophage lambda stabilization by auxiliary protein gpD: timing, location, and mechanism of attachment determined by cryo-EM. Structure 16, 1399–1406 (2008).

49. P. Singh, E. Nakatani, D. R. Goodlett, C. E. Catalano, A pseudo-atomic model for the capsid shell of bacteriophage lambda using chemical cross-linking/mass spectrometry and molecular modeling. J Mol Biol 425, 3378–3388 (2013).

50. E. Beghetto, N. Gargano, Lambda-display: a powerful tool for antigen discovery. Molecules 16, 3089–3105 (2011).

51. J. Nicastro et al., Construction and analysis of a genetically tuneable lytic phage display system. Appl Microbiol Biotechnol 97, 7791–7804 (2013).

52. L. Aghebati-Maleki et al., Phage display as a promising approach for vaccine development. J Biomed Sci 23, 66 (2016).

53. E. Jonczyk-Matysiak et al., Phage-Phagocyte Interactions and Their Implications for Phage Application as Therapeutics. Viruses 9, (2017).

54. A. Jegerlehner et al., A molecular assembly system that renders antigens of choice highly repetitive for induction of protective B cell responses. Vaccine 20, 3104–3112 (2002).

55. S. R. Leist et al., A Mouse-Adapted SARS-CoV-2 Induces Acute Lung Injury and Mortality in Standard Laboratory Mice. Cell 183, 1070–1085 e1012 (2020).

56. C. M. Coleman et al., MERS-CoV spike nanoparticles protect mice from MERS-CoV infection. Vaccine 35, 1586–1589 (2017).

57. J. Zhao et al., Rapid generation of a mouse model for Middle East respiratory syndrome. Proc Natl Acad Sci U S A 111, 4970–4975 (2014).

58. L. Piccoli et al., Mapping Neutralizing and Immunodominant Sites on the SARS-CoV-2 Spike Receptor-Binding Domain by Structure-Guided High-Resolution Serology. Cell 183, 1024–1042 e1021 (2020).

59. D. Corti et al., Prophylactic and postexposure efficacy of a potent human monoclonal antibody against MERS coronavirus. Proc Natl Acad Sci U S A 112, 10473–10478 (2015).

60. B. Rockx et al., Structural basis for potent cross-neutralizing human monoclonal antibody protection against lethal human and zoonotic severe acute respiratory syndrome coronavirus challenge. J Virol 82, 3220–3235 (2008).

61. W. B. Alsoussi et al., A Potently Neutralizing Antibody Protects Mice against SARS-CoV-2 Infection. J Immunol 205, 915–922 (2020).

62. M. A. Tortorici et al., Ultrapotent human antibodies protect against SARS-CoV-2 challenge via multiple mechanisms. Science 370, 950–957 (2020).

63. Y. Wu et al., A noncompeting pair of human neutralizing antibodies block COVID-19 virus binding to its receptor ACE2. Science 368, 1274–1278 (2020).

64. R. E. Chen et al., In vivo monoclonal antibody efficacy against SARS-CoV-2 variant strains. Nature, (2021).

65. Y. Guo et al., A SARS-CoV-2 neutralizing antibody with extensive Spike binding coverage and modified for optimal therapeutic outcomes. Nat Commun 12, 2623 (2021).

66. C. Kim et al., A therapeutic neutralizing antibody targeting receptor binding domain of SARS-CoV-2 spike protein. Nat Commun 12, 288 (2021).

67. A. Baum et al., REGN-COV2 antibodies prevent and treat SARS-CoV-2 infection in rhesus macaques and hamsters. Science 370, 1110–1115 (2020).

68. R. Shi et al., A human neutralizing antibody targets the receptor-binding site of SARS-CoV-2. Nature 584, 120–124 (2020).

69. A. Z. Wec et al., Broad neutralization of SARS-related viruses by human monoclonal antibodies. Science 369, 731–736 (2020).

70. K. O. Saunders et al., Neutralizing antibody vaccine for pandemic and pre-emergent coronaviruses. Nature 594, 553–559 (2021).

71. V. Manolova et al., Nanoparticles target distinct dendritic cell populations according to their size. Eur J Immunol 38, 1404–1413 (2008).

72. K. D. Brune, M. Howarth, New Routes and Opportunities for Modular Construction of Particulate Vaccines: Stick, Click, and Glue. Front Immunol 9, 1432 (2018).

73. T. K. Tan et al., A COVID-19 vaccine candidate using SpyCatcher multimerization of the SARS-CoV-2 spike protein receptor-binding domain induces potent neutralising antibody responses. Nat Commun 12, 542 (2021).

74. A. A. Cohen et al., Mosaic nanoparticles elicit cross-reactive immune responses to zoonotic coronaviruses in mice. Science 371, 735–741 (2021).

75. L. He et al., Single-component, self-assembling, protein nanoparticles presenting the receptor binding domain and stabilized spike as SARS-CoV-2 vaccine candidates. Sci Adv 7, (2021).

76. P. T. Heath et al., Safety and Efficacy of NVX-CoV2373 Covid-19 Vaccine. N Engl J Med, (2021).

77. S. Chiba et al., Multivalent nanoparticle-based vaccines protect hamsters against SARS-CoV-2 after a single immunization. Commun Biol 4, 597 (2021).

78. A. C. Walls et al., Elicitation of Potent Neutralizing Antibody Responses by Designed Protein Nanoparticle Vaccines for SARS-CoV-2. Cell 183, 1367–1382 e1317 (2020).

79. J. ter Meulen et al., Human monoclonal antibody combination against SARS coronavirus: synergy and coverage of escape mutants. PLoS Med 3, e237 (2006).

80. M. Yuan et al., A highly conserved cryptic epitope in the receptor binding domains of SARS-CoV-2 and SARS-CoV. Science 368, 630–633 (2020).

81. B. S. Graham, Rapid COVID-19 vaccine development. Science 368, 945–946 (2020).

82. F. Amanat et al., A serological assay to detect SARS-CoV-2 seroconversion in humans. Nat Med 26, 1033–1036 (2020).

83. D. F. Robbiani et al., Convergent antibody responses to SARS-CoV-2 in convalescent individuals. Nature 584, 437–442 (2020).

84. D. Pinto et al., Cross-neutralization of SARS-CoV-2 by a human monoclonal SARS-CoV antibody. Nature 583, 290–295 (2020).

85. J. Schindelin et al., Fiji: an open-source platform for biological-image analysis. Nat Methods 9, 676–682 (2012).

86. C. M. Coleman et al., Purified coronavirus spike protein nanoparticles induce coronavirus neutralizing antibodies in mice. Vaccine 32, 3169–3174 (2014).

87. V. M. Corman et al., Detection of 2019 novel coronavirus (2019-nCoV) by real-time RT-PCR. Euro Surveill 25, (2020).

