## Supplemental Fig S1 and Fig S2 for "Phage-like particle vaccines are highly immunogenic and protect against pathogenic coronavirus infection and disease"

### Figure S1

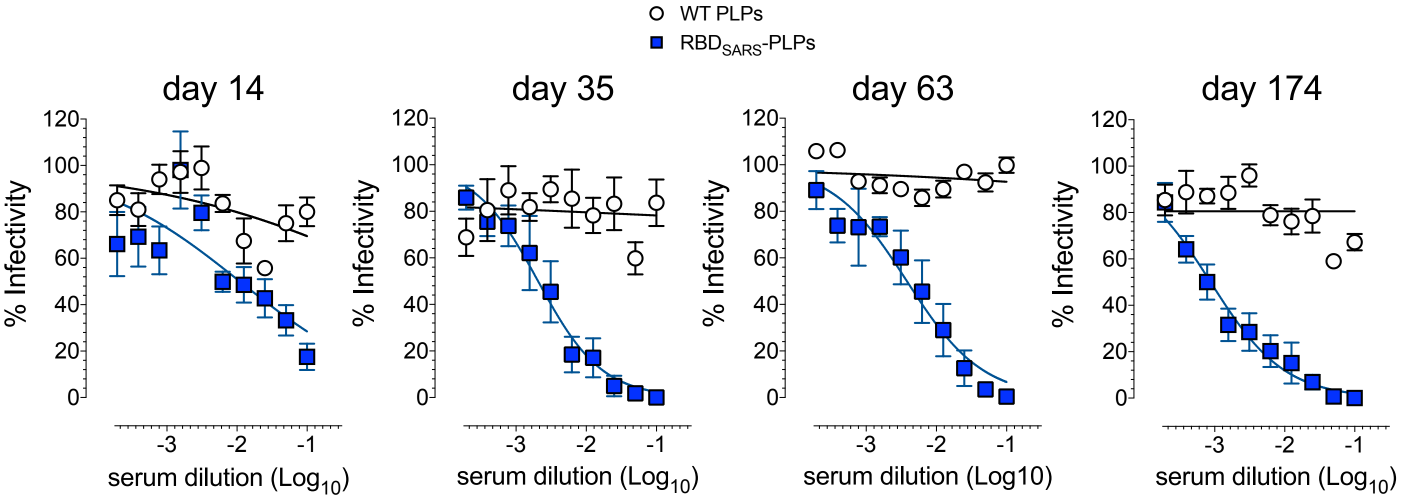


**Figure S1. Immunization with RBD$_{SARS}$-PLPs elicits potent and durable SARS-CoV-2 neutralizing antibodies.** WT BALB/c mice (n = 5/group) were immunized with 10 µg of WT PLPs (control) or 60% RBD$_{SARS}$-PLPs by intramuscular (i.m.) injection on days 0 and 21. Animals were bled on days 14, 35, 63, and 174 and SARS-CoV-2 neutralizing activity was determined by a focus reduction neutralization test (FRNT).

### Figure S2

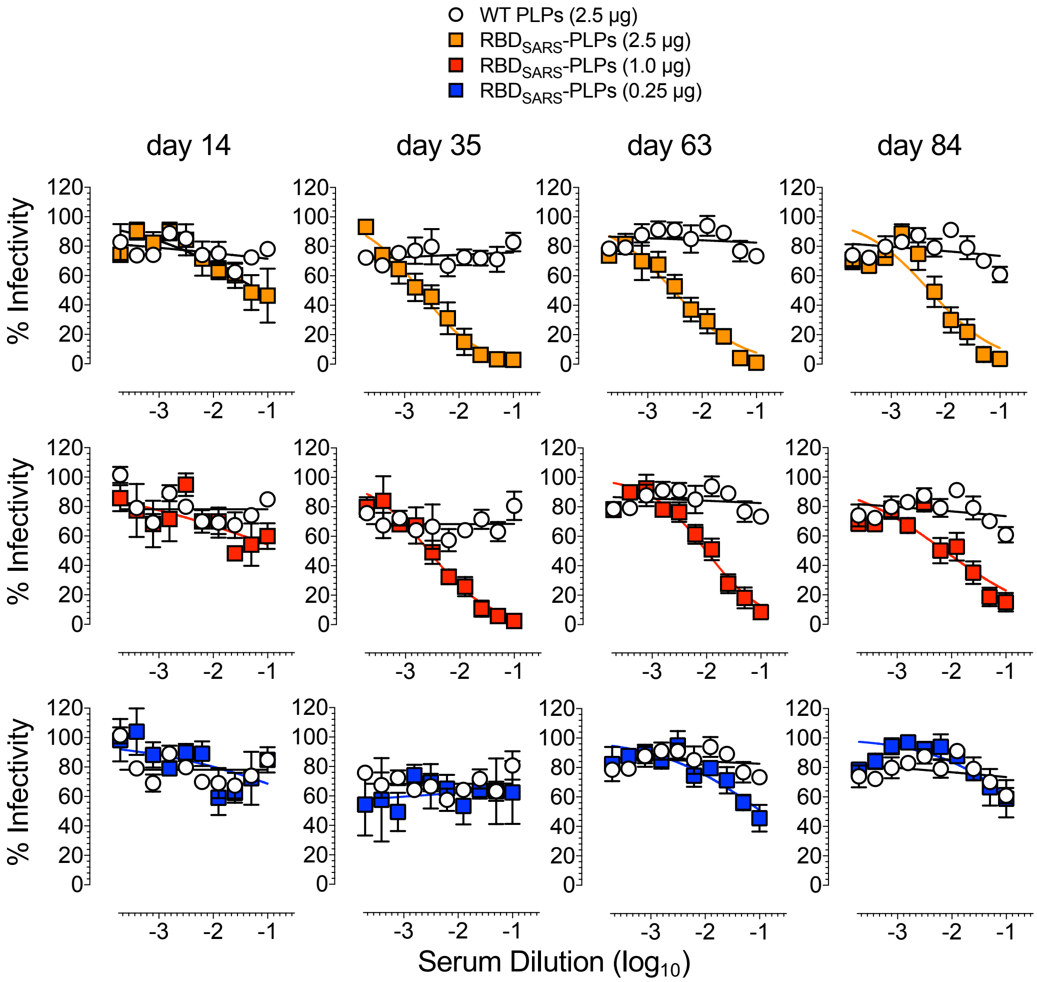


**Figure S2. Low dose immunization with RBD$_{SARS}$-PLPs elicits potent and durable SARS-CoV-2 antibody responses.** WT BALB/c mice (n = 5/group) were immunized with 2.5 μg of WT PLPs (control) or 2.5, 1.0, or 0.25 μg of RBD$_{SARS}$-PLPs by intramuscular (i.m.) injection on days 0 and 21. Animals were bled on days 14, 35, 63, and 84. SARS-CoV-2 neutralizing activity in serum samples was determined by a focus reduction neutralization test (FRNT).
